# Preclinical Proof of Concept for a personalized SNAP^TM^-TIL (Specific Neo-Antigen Peptides -TIL) therapy platform

**DOI:** 10.1101/2024.12.12.628191

**Authors:** Tithi Ghosh Halder, Erin Kelley, Jorge Soria-Bustos, Trason Thode, Serina Ng, Taylor Bargenquast, Alexis Weston, Ryan Rodriguez del Villar, Mohan Kaadige, Erkut Borazanci, Michael Gordon, Justin Moser, Frank Tsai, Saul J. Priceman, Stephen J. Forman, Yan Xing, John A. Altin, Raffaella Soldi, Sunil Sharma

## Abstract

Tumor-infiltrating lymphocyte (TIL) therapy, which involves extracting, expanding, and reinfusing immune cells to target cancer cells, has shown promise in melanoma treatment, but requires optimization for broader efficacy. The success of TIL therapy depends on the recognition of tumor-associated antigens, but neoantigen-reactive T-cells are often rare and exhausted in less immunogenic malignancies. Isolating T cells enriched in neoantigen reactivity prior to *in vitro* expansion and reinfusion may improve the response rates. To this end, our proprietary Specific Neo-Antigen Peptides (SNAP™) technology platform improves the accuracy of neoantigen prediction and validation by combining advanced computational modeling and PepSeq, a high-throughput screen for the physical credentialing of putative neoantigens based on their affinity to bind patient-specific HLA class II proteins. This approach allows for the education and enrichment of TILs (SNAP-TILs) with personalized, predefined, highly immunogenic neoantigens prior to expansion. Using the SNAP platform, we consistently achieved, on average, an SNAP-TIL product comprising 96% CD3+ cells, with a mixture of 75% effector and 23% central memory cells. SNAP-TILs exhibited greater efficacy and selectivity in immune infiltration than TIL, which was expanded by the rapid expansion protocol alone using *ex vivo* models. SNAP-TIL was also reactive in highly and poorly immunogenic tumors, with 70% and 50% tumor growth inhibition in melanoma and pancreatic patient-derived xenograft models, respectively. This study demonstrates the novel benefit of our Personalized Neoantigen Pipeline approach, potentially providing a durable antitumor immune response for a larger proportion of cancer patients.

## INTRODUCTION

The development of adoptive cell therapies (ACT) in solid tumors has lagged behind hematologic cancers for many reasons, including tumor heterogeneity, paucity of tumor-specific neoantigens, and possessing an immunosuppressive tumor microenvironment (TME). Unlike CAR T or other cell therapies, adoptive tumor-infiltrating lymphocyte (TIL) therapy has achieved promising results in solid tumors. It has sometimes generated complete responses in patients who have failed first-line therapies (1–3). Recently, the FDA approved Iovance’s Lifileucel, an unmodified but *in vitro* expanded TIL, to treat unresectable or metastatic melanoma. TIL therapy has distinct advantages over other cell therapies in treating solid tumors. TILs are polyclonal cells with multiple T cell receptors (TCR) derived directly from the tumor and can recognize a wide range of tumor-associated antigens (TAA), whereas CAR T cells require genetic engineering to recognize one or occasionally two neoantigens on tumor cells (4). Due to tumor heterogeneity and the expression of single neo-epitopes in normal cells, it is still doubtful that a CAR T approach directed against one or a few neoantigens is likely to succeed. TIL therapy, on the other hand, can target multiple neoantigens, overcoming the tumor heterogeneity conundrum and reducing the likelihood of tumor escape, particularly in solid tumors. Likewise, TILs can target unknown TAA within the TME, facilitating their migration toward the TME after infusion (5).

TIL therapy involves the expansion of a heterogeneous population of endogenous T cells found in an excised tumor, and its success depends largely on the recognition of specific TAAs, especially neoantigens. Consequently, TILs comprise effector memory T cells with high proliferation efficiency, antitumor functions, and chemokine receptors that may allow certain inhibitory immune regulators to bypass (6). Another advantage is that TILs are autologous products from the patient without gene modification, thus simplifying the manufacturing process (7). Most side effects stem from the associated lymphodepleting chemotherapy regimens or high-dose IL-2, but they are manageable (8). It also appears to be a long-lasting therapy with evidence of TILs patrolling years after a single infusion (3,9), possibly providing long-term protection against disease resurgence and progression.

The apparent advantages of TIL therapy are promising and suggest that this approach may be more effective against tumor cells than other ACT strategies. Despite the success of TIL treatment in cancers such as metastatic melanoma and cervical cancer (9–11), response rates in other types of solid tumors — such as colorectal cancer (CRC), cholangiocarcinoma, non-small cell lung cancer (NSCLC), head and neck cancers, and breast cancer — remain limited (5–7). This limitation stems from the fact that most solid tumors express few TAAs and thus are considered less immunogenic, making them less susceptible to immunotherapy (12–14). Neoantigen-reactive T cells in these cancers are often rare and exhausted, and recent studies have shown that *ex vivo* expansion can further diminish their effectiveness, complicating the development of highly reactive TIL products for treatment (8).

Having been isolated from tumors, TILs already recognize many neoantigens in cancer cells. However, many studies suggest that employing isolated T cells enriched in neoantigen reactivity prior to *in vitro* expansion and reinfusion may improve the response rate to TIL therapy in solid tumors (8,15–17). With deep-sequencing technologies, it has become feasible to identify the mutations present in the exome for an individual tumor with relative ease and thereby predict potential neoantigens. Neoantigen discovery and selection depend heavily on *in silico* prediction of neoantigenic epitopes/MHC affinity. However, only a small proportion of neoantigens selected by next-generation sequencing and *in silico* prediction algorithms induce an immune response in preclinical models and translational studies (18–20).

Relying only on *in silico* prediction of neoantigens could miss critically important immune responses (18–20), particularly in the case of immunologically cold tumors. Thus, we established a comprehensive ‘Personalized Neoantigen Pipeline’ to support personalized TIL therapy development. Our approach combines a highly multiplexed MHC class II binding assay based on libraries of DNA-barcoded peptides [“PepSeq” (21)] with *in-silico* prediction for MHC class I high-affinity peptides. Together, this new neoantigen identification technology improves neoantigen prediction to generate autologous TILs enriched for T cells that specifically recognize clonal neoantigenic epitopes we have termed ‘Specific Neo-Antigen Peptides’-TILs or SNAP™-TIL. Here, we evaluate the feasibility and efficacy of our SNAP-TIL product in cancer treatment using excised solid tumors from a cohort of 15 patients with various tumor types, including CRC, melanoma, uveal melanoma, NSCLC, and pancreatic ductal adenocarcinoma (PDAC).

## MATERIAL AND METHODS

### Patient biospecimens

Primary tumor samples, adjacent normal tissue, and peripheral blood samples were obtained from patients with stage III or IV disease who had received several lines of chemotherapy and underwent standard-of-care surgical resection. Cancer indications included CRC, melanoma, uveal melanoma, NSCLC, and PDAC. Blood samples were collected into an EDTA tube. Ficoll density gradient centrifugation was performed to isolate the peripheral blood mononuclear cells (PBMC). Isolated PBMC samples were cryopreserved with 90% FBS and 10% DMSO freezing media. Fresh tissue samples were dissected surgically, and tumor regions and normal adjacent were defined macroscopically and collected in MACS® Tissue Storage Solution (Miltenyi). Single-cell suspensions were prepared by mechanical and enzymatic tissue dissociation using the Human Tumor Dissociation Kit (Miltenyi) and then filtered, washed, and resuspended in RPMI-1640. CD45+TILs were isolated from the tumor digests using CD45 microbeads and maintained in RPMI-1640 (Sigma-Aldrich) modified with 10% human serum, penicillin-streptomycin, sodium pyruvate, non-essential amino acids, L-glutamine, and 2- mercaptoethanol. This study was approved by the Institutional Review Board of the HonorHealth Research Institute (IRB approval no. 1249216). All participants provided written informed consent before participating in the study.

### Neoantigen identification and selection

Neoantigens were identified by whole exome sequencing (WES) of tumor and germline DNA from peripheral blood and then selected based on these properties: nonsynonymous point mutations, frameshift mutations, gene fusions, allele-specific expression of the mutation, tumor clonal epitopes, oncogenes, tumor mutation burden, and peptide manufacturing scores. The patient’s HLA haplotypes were determined by clinical grade typing using conventional locus-specific primers and dideoxy-based Sanger sequencing to ensure typing accuracy (22), as inferred from WES. Following the “tumor versus normal” WES, a pool of oligonucleotides encoding peptides of interest was synthesized and used to produce a library of peptide:DNA conjugates [using the PepSeq platform (21)] to empirically identify binders to patient-specific MHC class II proteins as described below. In parallel, an *in-silico* predictive algorithm (netMHCpan) was used to identify binders to MHC class I. Peptides with a minimum predicted MHC binding percentile rank (lower rank corresponds to stronger binding) regarding a random set of peptides were selected to include in TIL:peptide co-culture. Predictions were generated for each patient’s MHC class I alleles (up to six). A custom pool of ∼30 peptides with roughly an equal mix of the best class I and class II binders was selected for each patient and utilized to ‘educate’ TILs.

### PepSeq assay

PepSeq libraries were generated as previously described (21). We designed 3-tiled 15-mer peptides for each tumor mutation, with mutant residues in positions 7, 8, and 9. We then encoded these into a patient-specific oligonucleotide library of up to 7,500 peptide sequences. This library was converted into a DNA-barcoded peptide library using 4 *in vitro* enzymatic steps: transcription, ligation to a puromycin adapter oligo, translation (including intramolecular puromycin coupling), and reverse transcription (21). We then used these libraries to assess binding to purified HLA proteins (23,24). Specifically, purified CLIP-tethered HLA-DR protein(s) matching the patient’s genotype was sourced from the NIH Tetramer Core Facility, and the CLIP peptide was detached by cleavage using the relevant protease (thrombin or 3C). 0.5μg of purified recombinant HLA-DR was then incubated overnight with 1pmol PepSeq library in binding buffer (2.3% n-octylglucoside, 230mM NaCl, 110mM citrate, 4.7mM EDTA, 1.3% protease inhibitor cocktail) at pH 5.5. Reactions without MHC or thrombin cleavage were used as binding controls. Next, protein G magnetic Dynabeads (ThermoFisher) pre-incubated with 0.5mg/mL anti-HLA-DR clone L243 (BioLegend) was used to immunoprecipitate the MHC/PepSeq library pools for 2h at RT. Beads were washed 10 times to remove DNA-barcoded peptides not attached to bead-bound MHC. The PepSeq eluate sequencing and quantification of peptide-level enrichment were performed as previously described (21). In brief, raw sequence reads from the PepSeq library were analyzed against reference sequences to produce a matrix of read counts, which were then normalized and compared to controls to select peptides for TIL co-culture experiments based on a constant-adjusted binding score ratio.

### SNAP-TIL generation

The selected neoantigen peptides were co-cultured with the harvested TILs to obtain TILs selective for the patient’s tumor neoantigens (SNAP-TIL). First, TILs were isolated from the single-cell suspension of the resected tumor using CD45+ microbeads. Next, isolated TILs were incubated for 7 days with a selected pool of high-ranking neopeptides. After incubation, a peptide-enriched TIL population was generated. Finally, the TILs were expanded with the rapid expansion protocol (REP) using IL-2 and anti-CD3 (Miltenyi) in the presence of irradiated feeder lymphocytes into 40mL capacity flasks with a 10cm² gas-permeable silicone bottom (G-Rex10, Wilson Wolf) as previously described (25).

### Single-cell sequencing

Cryopreserved TILs cultured with neopeptides were thawed, washed, and resuspended in PBS with BSA and RNase inhibitors. TILs were assessed for viability using Calcein-AM and NucBlue Live ReadyProbes Reagent, then sorted using the FITC channel on the Sony SH800S. Sorted cells were processed using the 10x Genomics Chromium Next GEM Single Cell 5’ v2 Dual Index kit, amplified with cDNA libraries constructed per protocol. Quality control was conducted using Agilent Tapestation 4200 HS D1000 and Kapa Library Quantification Kit, followed by sequencing on Illumina’s iSeq 100 v2 and NovaSeq 6000 S4 v1.5 platforms.

### Immune infiltration assay

Tumor spheroids (i.e., tumoroids) were generated from patient-derived primary tumor cells using a protocol previously published (26). Briefly, TILs were labeled with Molecular Probes Vybrant CM-Dil Cell Labeling Solution (RFP; ThermoFisher). Next, cells were washed with PBS twice and resuspended in the appropriate complete growth medium. A 5μm HTS Transwell 96- Well Permeable Support receiver plate (Corning) was placed on each ultra-low attachment spheroid microplate to allow SNAP-TIL infiltration into the spheroids. Molecular Probes Vybrant CM-Dil-stained TILs were then seeded into inserts at 5×10^5^Lcells/well to ensure a tumoroid:TIL ratio of 1:10. After 24h, inserts were removed, and tumoroid microplates were analyzed by 3D Z-stack imaging and morphometric analysis with Cytation 5 software.

### IFN**γ** ELISpot assay

The activation of SNAP-TIL by individual neopeptides was measured using an interferon-gamma (IFN-γ) ELISpot assay. Briefly, multi-screen plates were coated with capture antibodies for human IFNγ. 2×10^5^ SNAP-TIL cells/well were co-incubated for 24h with each neopeptide (2.5μg). Plates were washed, and biotinylated anti-IFNγ was added, followed by streptavidin conjugated-alkaline phosphatase. Spots were developed with BCIP/NBT. Each plate included positive (1ng/ml PMA and 500ng/ml Ionomycin) and negative (DMSO) controls co-incubated with the peptide pool to confirm reagent performance. Spots were imaged and enumerated using an Immunospot analyzer (Bioreader 7000). Responses were positive if the spot-forming cell count detected was ≥10 spots over control, +3 standard deviations, and confirmed in repeat experiments.

### Flow cytometry FACS and immunophenotyping

Single-cell tumor suspensions were prepared through mechanical and enzymatic dissociation using a Milteyni MACs tissue dissociator. Next, for surface staining, cells were washed, counted, and dissolved in Cell Staining buffer (Biolegend) and incubated at 4°C for 15min with fluorochrome-conjugated antibodies. Cells were permeabilized and stained with anti-human FOXP3 (Biolegend) for intracellular staining of Tregs per the manufacturer’s instructions. BV421+ and single cells were gated out, followed by CD45+ selection. All flow cytometry FACS analyses were performed on a Sony SH800, and the data was analyzed using FlowJo (RRID:SCR_008520) (BD Life Sciences). Supplementary Table S1 lists all antibodies used for flow cytometry.

### qPCR

Total RNA was isolated using the RNeasy Mini Kit (Qiagen) and quantified by spectroscopy (Nanodrop ND-8000). Samples were then reverse transcribed to cDNA using a high-capacity cDNA reverse transcription kit. cDNA was amplified, detected, and quantified using SYBR green reagents and the ViiA 7 Real-Time PCR System (Applied Biosystems). Data were normalized to GAPDH expression. A list of primers is in Supplementary Table S2.

### Measurement of Granzyme B and IFN**γ**

Meso Scale Discovery (MSD) technology platform was used to measure granzyme B and IFN- γ secretion from TILs co-cultured with autologous tumor cells. Briefly, TIL cells were placed into a 24-well plate with autologous tumor cells in a 1:1 ratio. After a 24h incubation, supernatants were harvested, and granzyme B and IFN-γ release were quantified by the MSD assay, following the manufacturer’s instructions.

### Patient-derived xenograft (PDX) models

Excised tumors from melanoma or PDAC patients were mechanically dissociated, filtered through a cell strainer, washed, and resuspended in RPMI-1640. PDX tumors were then propagated by subcutaneously implanting 5×10^6^ cells mixed with an equal volume of matrigel (Corning) into the flank of 8-week-old female NOG mice (Taconic Biosciences). Tumors were extracted at ∼2000mm^3^ and then dissociated as previously described. Next, 2×10^6^ cells were subcutaneously implanted into human *IL-2* transgenic NOG (*NOD.Cg-Prkdcscid Il2rgtm1Sug Tg(CMV-IL2)4-2Jic/JicTac*, Taconic). Animals were randomly divided into treatment groups (n=10 per group) of equal proportions of tumor sizes (∼200mm^3^), and 1.8×10^7^ autologous TILs were administered by retroorbital injection. Tumor growth was measured thrice per week using calipers, and tumor volume was calculated: (length × width^2^)/2. Once tumor volumes surpassed 2000mm^3^, mice were humanely sacrificed, and tissue was collected for *ex vivo* analysis. All animal experiments were approved by the University of Arizona Institutional Animal Care and Use Committee (IACUC approval no. 18-494).

### Immunohistochemistry (IHC)

PDX tumors were fixed in a 10% neutral buffered formalin solution overnight and then transferred to 70% ethanol before being embedded in paraffin blocks. Serial cryostat tissue sections (4µm) were mounted onto positively charged glass slides, deparaffinized, and rehydrated. Antigen retrieval was performed using Diva Decloaker and immersed in a blocking reagent for 10min. DAB chromogen staining (Cell Signaling) was added after incubation with secondary antibodies. The following Abcam antibodies were used: anti-CD3 epsilon (ab16669), anti-CD4 (ab213215), and anti-CD8 alpha (ab237709). Slides were counterstained with hematoxylin, mounted with Pertex medium, and imaged on an AperioAT2 Scanner (Leica Biosystems).

### Statistical analysis

For ELISpot and cytokine panel, data was log-transformed and analyzed by one-way ANOVA followed by the Dunnett or Tukey test for multiple comparisons. For the quantification of antigen-specific T cells and regulatory cells per field by cell counting, a paired t-test on log-transformed data was used. An unpaired 2-tailed student t-test was used to determine statistical significance between the different groups (e.g., responders vs. non-responders or serial time points). For IHC data, associations between two categorical variables were assessed using Fisher’s exact test and between categorical and continuous variables using Wilcoxon rank-sum or Kruskal– Wallis tests. The probability of survival was analyzed by Kaplan-Meier survival curves using the log-rank (Mantel-Cox) test. All data were screened for parametric statistical test assumptions. All statistical tests were set at the a priori alpha level of 0.05.

## RESULTS

### SNAP-TIL workflow

We have developed a new TIL therapy workflow (**Fig. 1A**) with improved neoantigen prediction to generate autologous TILs enriched with T cells *ex vivo* that specifically recognize clonal neoantigenic epitopes which we termed SNAP-TIL. Our approach combines a platform for highly multiplexed production and assay of customizable peptide sets [PepSeq (21)], a scalable proteomics platform for neoantigen discovery, with the *in-silico* prediction for MHC class I high-affinity peptides. Representative outputs from the PepSeq assay identifying HLA-binding peptides corresponding to tumor mutations are shown in **Fig. 1B** for patients with CRC, melanoma, or breast cancer. Briefly, TILs were isolated from single-cell suspension prepared from a resected tumor using CD45+ selection. Next, isolated TILs were incubated for 7 days with the selected custom pool of high-ranking neopeptides. After incubation, a peptide-enriched TIL population (SNAP-TIL) was generated and expanded by REP using IL-2 and anti-CD3 stimulation on irradiated autologous PBMC (27,28). Overall, the SNAP-TIL platform effectively enhances the ability of the immune system to attack the tumor cells in the body, as clonal neoantigens, unlike TAAs, are present only in tumor cells and are the most relevant therapeutic targets for T cells.

**Figure 1.**
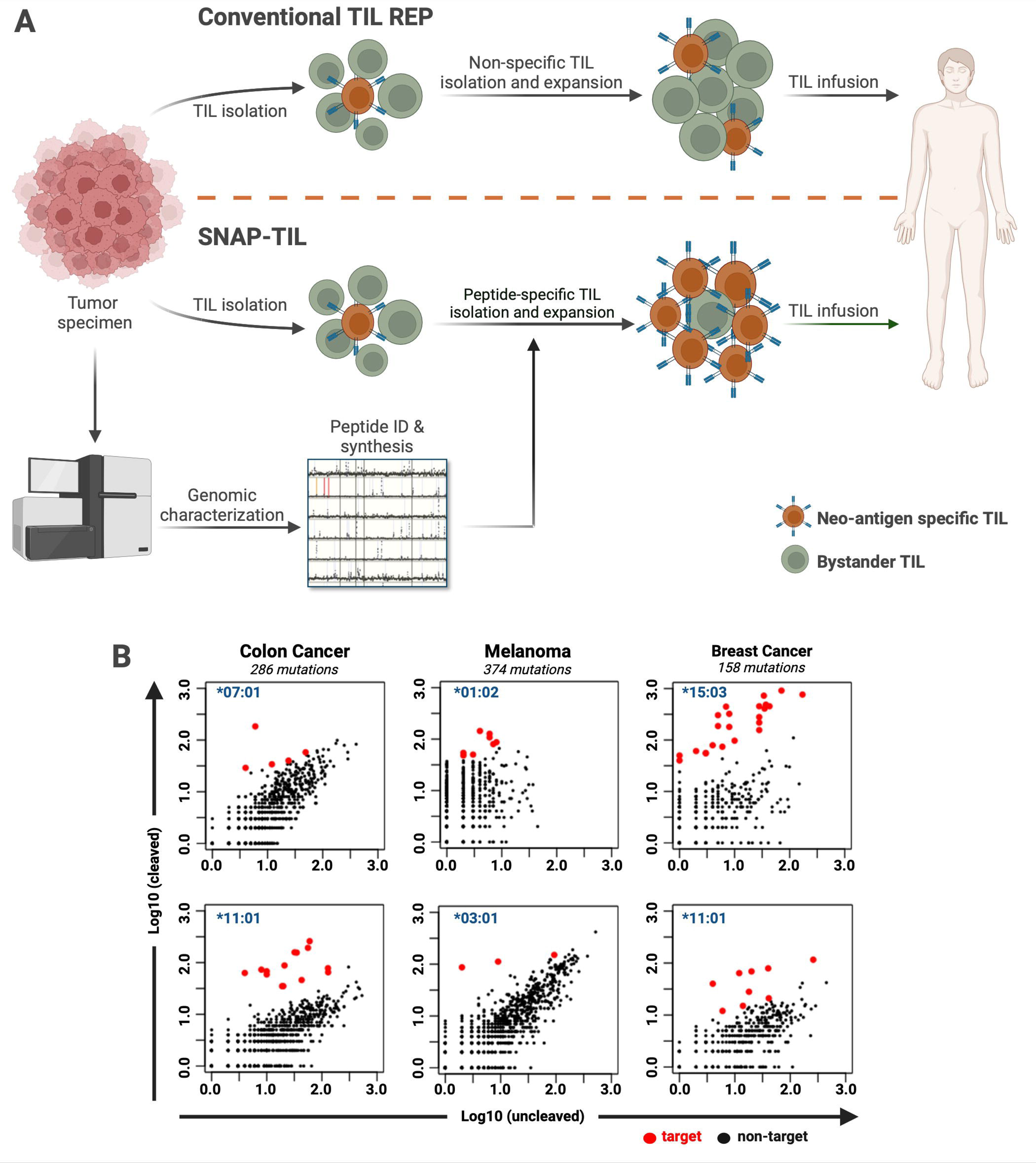
Overview of the SNAP-TIL platform to generate personalized, neoantigen-educated TILs. (A) Schematic workflow of the SNAP-TIL platform compared to the conventional rapid expansion approach (TIL REP). Identifying candidate neoantigens for a cancer patient begins with whole exome sequencing (WES, or whole genome) of excised tumor and germline DNA from peripheral blood to pinpoint tumor-specific mutations. The patient’s human leukocyte antigen (HLA) haplotypes are determined by clinical grade typing and inferred from WES. *In-silico* predictions combined with PepSeq are performed to identify the highest affinity binding neoantigens. A ranked list of up to 30 neopeptides is used to stimulate the expansion of neoantigen-specific T cells. (B) Identification of HLA-binding peptides corresponding to tumor mutations. Representative outputs from the PepSeq assay for three patients with colorectal cancer (left panel), melanoma (middle panel), or breast cancer (right panel). The ‘target’ neopeptides (red dots) are differentiated from non-target peptides (black dots). The asterisk indicates HLA-DRB1 haplotype. Figure created with BioRender.com.

### Educated SNAP-TIL is enriched with cytotoxic and effector memory T cells

The distribution of immune cell types, including T, B, natural killer (NK), and myeloid cells, was evaluated in SNAP-TIL post-REP. Flow cytometry analysis of SNAP-TIL from a cohort of CRC, melanoma, uveal melanoma, NSCLC, and PDAC patients showed that, on average, >96% of the SNAP-TIL were CD3+ T cells, a mixture of CD4+ and CD8+ T cells (**Fig. 2A**, top panel; Supplementary Fig. S1A-B). NK and B cells were generally detected in minuscule amounts (<3%) or undetected (**Fig. 2A**, top panel; Supplementary Fig. S1C-D). SNAP-TIL showed higher levels of expression for CD28 co-stimulatory molecules on CD8+ T cells, an indicator of increased responsiveness to antigenic stimulation (Supplementary Fig. S1E) and <3% Treg cells (Supplementary Fig. S1F-G). SNAP-TIL comprised 75% effector memory cells (CD197-/CD45RA-) and 23% central memory cells (CD197+CD45RA-; **Fig. 2A**, bottom panel, Supplementary Fig. S1F-G). Further analysis identified a subset of memory T cells, the stem-like memory T cells (TSMC), critical for *in vivo* expansion, long-term persistence, and antitumor activity (29). These data were confirmed by single-cell sequencing (Supplementary Fig. S2). In addition, single-cell sequencing of a PDAC patient SNAP-TIL after REP, we observed high levels of cytotoxic (CCL5, GZMK, GNLY, KLRG, and NKG7, Supplementary Fig. S2F) and proliferative (LIF, NME1, FABP5, and ORC6, Supplementary Fig. S2G) markers. Tumor cells were not detected in the final SNAP-TIL product (Supplementary Fig. S1H).

**Figure 2.**
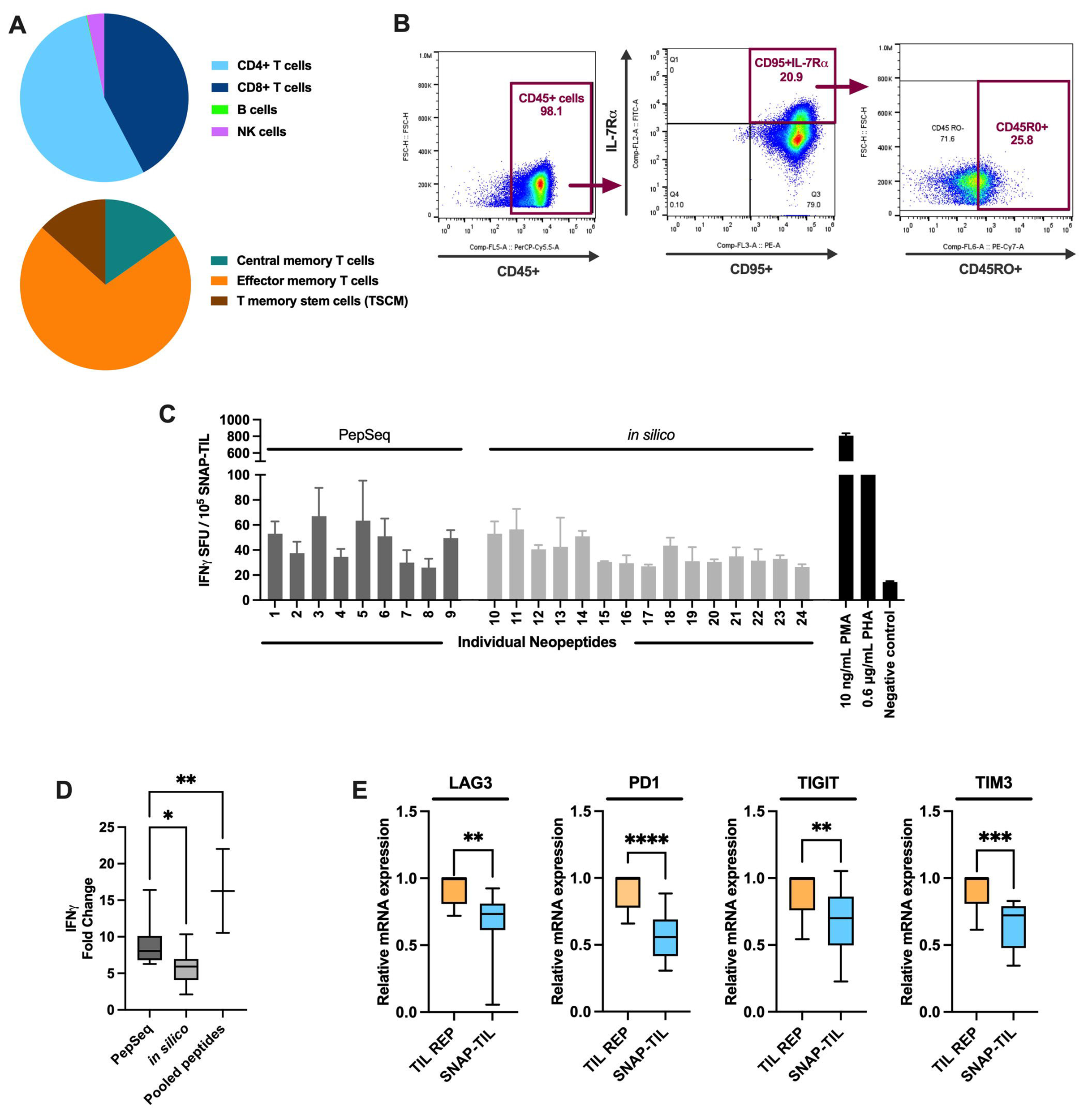
Immunophenotype of SNAP-TIL after rapid expansion. (A) Distribution of lymphocyte cell populations in SNAP-TIL from a compilation of tumor types (n = 3 to 6 patients). The top panel represents the percentile of CD4+ cells, CD8+ cells, B cells, and natural killer (NK) cells. The bottom panel represents the percentile of central memory, effector memory, and stem-like memory T cells (TSCM). (B) Flow cytometry analysis of SNAP-TIL for TSCM from a representative patient. Numbers indicate the percentage of cells in different gated populations. (C) Stimulation of SNAP-TIL from a PDAC patient in the presence of individual neopeptides measured by IFNγ ELISpot assay. PMA and PHA served as positive controls and DMSO as negative controls. Representative data from one of 3 independent experiments. Bars represent mean ± SD, n = 3 replicates, \**p* < 0.05, \*\**p* < 0.01, one-way ANOVA with Dunnett’s post-hoc test. (D) Measurement of IFNγ levels in SNAP-TIL expanded with PepSeq pool, *in silico* pool, or a combination of the two peptide pools. (E) TIL exhaustion was evaluated by qPCR of markers associated with replicative senescence and exhaustion of lymphocytes. In box-plot diagrams, the median is represented with a line, the interquartile range with a box, and the minimum and maximum of the data with the whiskers, n = 10 (5 tumor types), \*\**p* < 0.01, \*\*\**p* < 0.001, \*\*\*\**p* < 0.0001, one-way ANOVA with Dunnett’s post-hoc test.

The neoantigen-reactivity of SNAP-TIL was evaluated by the IFNγ ELISpot, which quantitatively measures the frequency of cytokine secretion from a single cell. PepSeq and *in-silico* peptides solicited an immune response from the SNAP-TIL (**Fig. 2C**; Supplementary Fig. S3A). However, when we compared the efficacy of the peptides selected by the PepSeq with that of the peptides selected by the *in-silico* algorithm, we observed that the PepSeq peptide pool produced higher levels of IFNγ, although the difference was not always statistically significant (**Fig. 2D**, Supplementary Fig. S3B-D). Notably, SNAP-TIL maintained functionality despite the 7-day incubation with the neoantigen peptides. We then evaluated TIL exhaustion status using gene expression markers associated with replicative senescence and exhaustion of lymphocytes, such as *LAG3*, *PD1*, *TIGIT*, and *TIM3*. qPCR analysis across this gene set showed that the SNAP-TIL product had a significantly reduced exhaustion status compared to TILs expanded by the rapid expansion protocol (REP) only, termed TIL REP (**Fig. 2E**).

### SNAP-TIL is robustly efficacious in patient-derived tumoroid models

SNAP-TIL efficacy was evaluated by the immune infiltration assay, a neoantigen immunogenicity assay we previously described (26). This assay involves generating and optimizing immune spheroids to test the ability of immune cells to interact with tumors derived from the same patient (26). This platform has the advantage of being reproducible and high throughput, and its effectiveness in evaluating immune response has been established. We evaluated tumoroids generated from CRC, melanoma, uveal melanoma, NSCLC, and PDAC patients. Briefly, fluorescently labeled SNAP-TIL and TIL REP were co-cultured with tumoroids prepared from biopsy tissue from the same patient in a transwell system to measure cellular infiltration and tumor reactivity. After 24h, the inserts were removed, and lymphocyte infiltration was quantified by 3D z-stack imaging and morphometric analysis. In this *ex vivo* model, SNAP- TIL demonstrated significantly higher efficacy in immune infiltration across all patient cohorts than naïve TIL (untreated) or TIL REP (**Fig. 3A**, Supplementary Fig. S4). We then examined the phenotype and function of SNAP-TIL. qPCR analysis revealed that SNAP-TIL had higher expression of lytic markers (granzyme B) and cytokine (IFNγ) markers after exposure to tumoroids (**Fig. 3B**, Supplementary Fig. S5). MSD analysis of conditioned medium from tumoroids and TIL co-cultures consistently revealed significantly high levels of granzyme B and IFNγ in SNAP-TIL (**Fig. 3C**, Supplementary Fig. S6). Together, these data show enhanced activation and cytotoxicity for SNAP-TIL compared to TIL REP.

**Figure 3.**
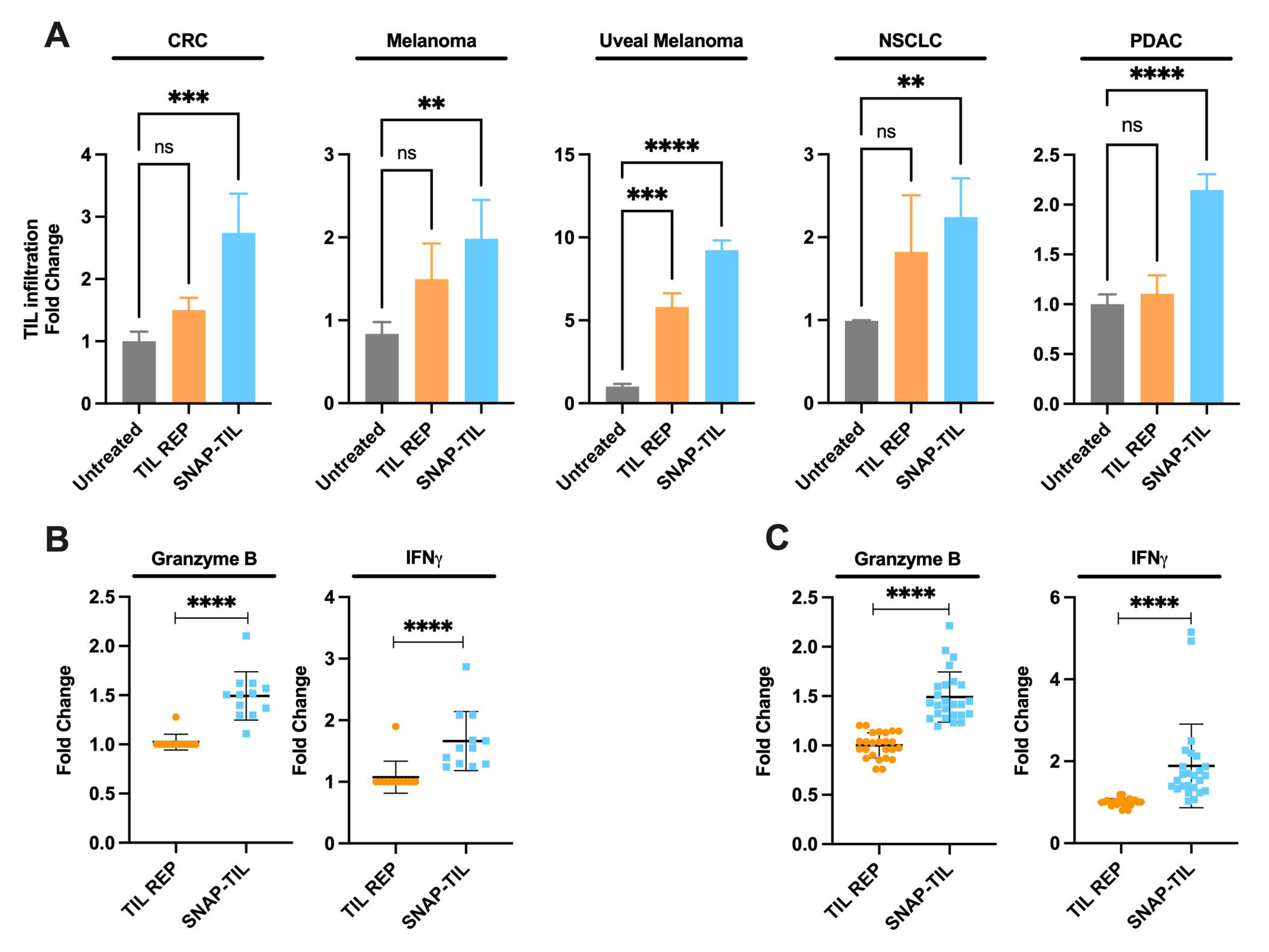
SNAP-TIL is superior to TIL obtained only by conventional rapid expansion protocol (TIL REP) in patient-derived spheroid models. (A) TIL infiltration into tumoroids after ‘educated’ with neopeptides (SNAP-TIL) is significantly higher than nonspecific, conventional TIL REP. Bar graphs represent mean ± SD, n = 2 - 6, \**p* < 0.05, \*\**p* < 0.01, \*\*\**p* < 0.001, \*\*\*\**p* < 0.0001, ns, not significant, ordinary one-way ANOVA with Tukey’s post-hoc test. (B) qPCR of granzyme B and IFNγ in TILs co-cultured with autologous *ex-vivo* tumoroids. n = 9 (C) Measurement of granzyme B and IFNγ levels in SNAP-TIL by MSD show cytotoxic properties (granzyme B; left panel) and enhanced activation (IFNγ; right panel). The horizontal black bars represent the mean ± SD. n = 9 - 11, ****p < 0.0001, unpaired t-test. CRC, colorectal cancer; NSCLC, non-small cell lung cancer; PDAC, pancreatic ductal adenocarcinoma

### *Ex vivo* SNAP-TIL responses are specific, selective, and durable

The specificity of SNAP-TIL-expanded T cell clones to autologous or allogeneic peptides was measured using the IFNγ ELISpot assay. SNAP-TIL generated from a PDAC patient was incubated with either a peptide pool prepared for the same patient or a peptide pool generated for a melanoma patient. SNAP-TIL reacted preferentially to autologous peptides, with no significant reaction to allogeneic peptides (**Fig. 4A**, left panel). Similar results were obtained using SNAP-TIL from a CRC patient incubated with allogeneic peptides for a PDAC patient (**Fig. 4A**, right panel). To confirm ELISpot assay reactivity using an orthogonal technique, we performed an *ex vivo* immune infiltration assay using patient-derived tumoroids. Briefly, melanoma patient-derived tumoroids were co-cultured with naïve TIL, TIL REP, or SNAP-TIL derived from a PDAC patient, and lymphocyte infiltration was quantified as previously described at 24, 48, and 72h. SNAP-TIL demonstrated no cross-reactivity to the allogeneic tumoroid, while TIL REP did (**Fig. 4B**).

**Figure 4.**
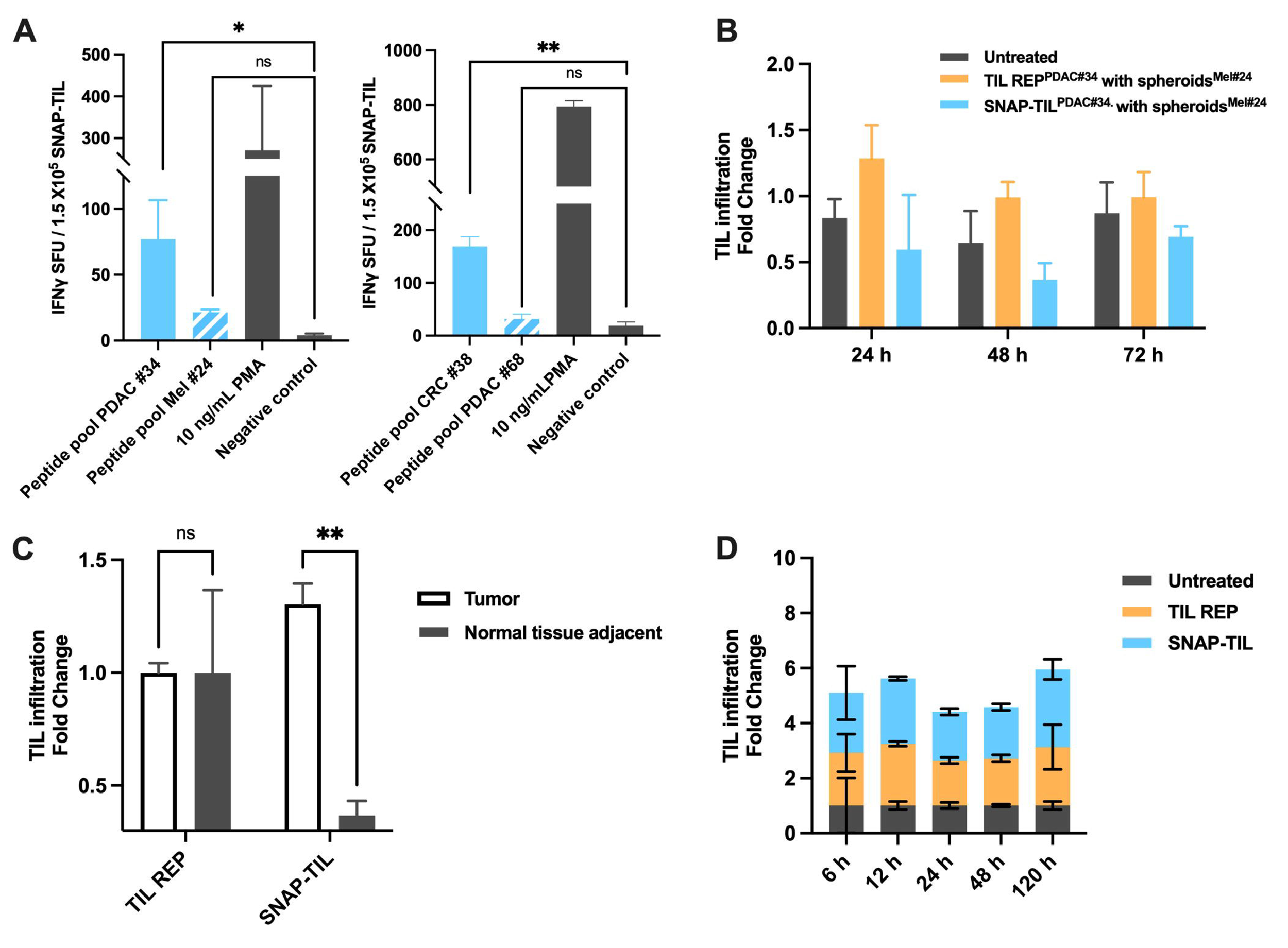
Immune response specificity of SNAP-TIL. (A) Immune reactivity measured by *ex-vivo* IFNγ ELISpot assay to allogeneic peptides versus autologous peptides. Representative response from PDAC patient #34 incubated with allogeneic peptides from melanoma patient #24 (left panel) and CRC patient #38 incubated with allogeneic peptides from PDAC patient #68 (right panel). Each bar represents the mean ± SD, n = 2, \**p* < 0.05, \*\**p* < 0.01, ns, not significant, ordinary one-way ANOVA with Tukey’s post-hoc test. (B) Immune reactivity by *ex-vivo* immune infiltration assay of allogeneic peptides at 24, 48, and 72h. Representative response from SNAP-TIL and conventional TIL REP from a PDAC patient incubated with allogeneic melanoma tumoroids. (C) A comparison of TIL infiltration in tumors and normal adjacent tissue from melanoma patients shows that SNAP-TIL selectively recognizes tumor tissue. Each bar represents the mean ± SD, n = 2 – 6 technical replicates, \**p* < 0.05, \*\**p* < 0.01, \*\*\**p* < 0.001, \*\*\*\**p* < 0.0001, two-way ANOVA with Šidák post-hoc test. (D) SNAP-TIL demonstrates higher immune reactivity that persists over days compared to TIL REP in melanoma tumoroids. Each bar represents the mean ± SD, n = 2.

One of the most striking toxicities specific to TIL therapy is those arising from a direct T-cell attack on normal tissues. This can take the form of an “on-target, off-tumor” autoimmunity (30). To investigate the potential of “off-target” autoimmunity, we evaluated SNAP-TIL selectivity toward a patient’s tumor tissue versus normal adjacent tissue (NAT) originating from the same patient. SNAP-TIL and TIL REP were co-cultured with the same patient-derived tumoroids or NAT spheroids to measure cellular infiltration and tumor reactivity. After 24h, lymphocyte infiltration was quantified by 3D z-stack imaging and morphometric analysis. *Ex vivo* responses showed SNAP-TIL favoring tumoroids over NAT, while no difference was observed in tumoroids derived from tumor and NAT treated with TIL REP (**Fig. 4C**). This suggests SNAP-TIL are enriched with T cells against ‘driver’ neoantigens and, thus, would expect to be associated with the lowest probability of antigen-escape relapses or autoimmunity when used in adoptive cell transfer protocols. We then tested the persistence of the SNAP-TIL immune response. Briefly, melanoma patient-derived tumoroids were co-cultured with naive TIL, TIL REP, or SNAP-TIL as previously described, and lymphocyte infiltration was quantified by 3D z-stack imaging and morphometric analysis at 6, 12, 24, 48, and 120h. SNAP-TIL showed significantly higher efficacy in immune infiltration, and the effect was maintained during the experiment (**Fig. 4D**). Altogether, these data demonstrate SNAP-TIL to be specific, selective, and durable.

### SNAP-TIL infiltrates and controls tumor growth in a melanoma PDX model

The *in vivo* efficacy of SNAP-TIL was examined in a melanoma PDX model using *hIL2*-NOG mice. Tumor-bearing mice were administered 1.8 × 10^7^ of SNAP-TIL or TIL REP or remained untreated as a control group. Tumor size was monitored weekly for 8 weeks. SNAP-TIL significantly delayed tumor and, by the study end, achieved 67.6% tumor growth inhibition (TGI) compared to 28.7% TGI with TIL REP (**Fig. 5A**). No discernable change in body weight was observed in the treatment groups (Supplementary Fig. S7A). Survival rates following SNAP-TIL treatment were significantly higher (*p*=0.0028) than the control group (**Fig. 5B**). Fold change in granzyme B and IFNγ levels measured by MSD were significantly higher in SNAP-TIL versus TIL REP or untreated mice (**Fig. 5C**). IHC analysis of comparable tumor area showed significantly higher numbers of CD4+ and CD8+ T cells in SNAP-TIL treated mice compared to TIL REP treated mice (**Fig. 5D**), confirming SNAP-TIL traffics to the tumor as we observed in the *ex vivo* tumoroid model (**Fig. 3A**). Further, we observed more than doubling CD8+ T cells in the spleen from SNAP-TIL versus TIL REP treated mice (**Fig. 5E**), suggesting that SNAP-TIL promotes immune homeostasis and CD8+ T cell expansion after infusion. We then examined the tumor lymphocyte distribution in SNAP-TIL-treated mice measured by flow cytometry. It showed that CD4+ and CD8+ T cells dominated with only a minor fraction of NK or B cells (**Fig. 5F**). Last, we measured neoantigen reactive T cell clones after SNAP-TIL treatment. CD45+ cells isolated from the melanoma PDX tumors were co-cultured with each neopeptide, and reactivity was measured using the IFNγ ELISpot assay. A total of 6 out of the 26 neopeptides (23%) produced a T-cell response of sufficiently high magnitude compared to the baseline (**Fig. 5G**), suggesting *in vivo* T-cell expansion.

**Figure 5.**
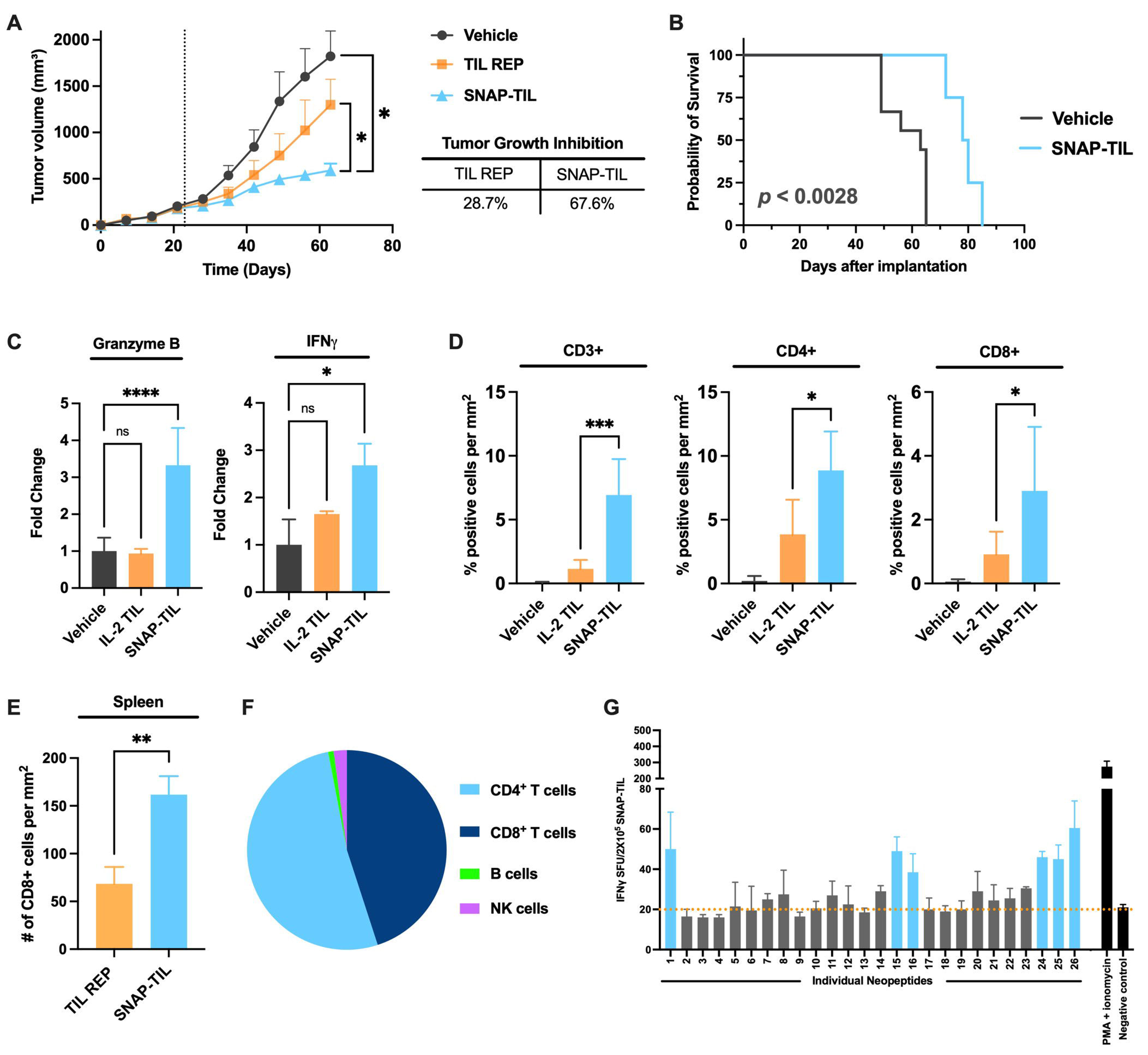
SNAP-TIL efficacy is superior to conventional TIL REP in a melanoma PDX model. (A) Female *hIL2*-NOG mice were subcutaneously implanted with propagated melanoma cells. Once tumors reached around 200 mm^3^, indicated by a dotted line, autologous SNAP-TIL, TIL REP, or vehicle was administered by retroorbital injection. Tumor volume was measured weekly and plotted as mean + SEM, n = 10 mice/group, \**p* < 0.05, unpaired 2-tailed student t-test. (B) Kaplan-Meier survival probability. (C) Fold change of Granzme B and IFNγ levels in TILs from PDX tumors measured by MSD assay. Each bar represents the mean ± SD, n = 6, \**p* < 0.05, \*\*\*\**p* < 0.0001, ns, not significant, ordinary one-way ANOVA with Dunnett’s post-hoc test. (D) Quantification of CD3+, CD4+, and CD8+ cells in melanoma PDX by IHC. Each bar represents the mean ± SD, n = 3, \**p* < 0.05, \*\*\**p* < 0.001, unpaired t-test. (E) Quantification of CD8+ T cells in spleen by IHC. Each bar represents the mean ± SD, n = 5, \*\**p* < 0.01, unpaired t-test. (F) Average distribution of CD4+ cells, CD8+ cells, B cells, and NK cells within the CD45+ population of SNAP-TIL in melanoma PDX tumors (n = 3) by flow cytometry. (G) Detection of neoantigen reactive T cell clones after SNAP-TIL treatment in the melanoma PDX model. CD45+ cells isolated from the melanoma PDX tumors were co-cultured with each neopeptide, and reactivity was measured using the IFNγ ELISpot assay. Each bar represents the mean ± SD. Reactive neopeptides are indicated in blue. The dotted orange line represents the baseline.

### SNAP-TIL demonstrates *in vivo* efficacy in an immunologically cold tumor

Given that the reliance on *in silico* prediction of neoantigens alone may miss critically important immune responses, particularly in the case of immunologically cold tumors, we sought to determine whether SNAP-TIL would outperform conventional TIL REP in a PDAC PDX model. We generated PDAC PDX *hIL2*-NOG mice and treated them as described above for melanoma. Tumor size was monitored weekly for 5 weeks. As seen with melanoma, SNAP-TIL significantly delayed tumor growth and, by the study end, achieved 51.8% TGI compared to 18.5% TGI with TIL REP (**Fig. 6A**). No discernable change in body weight was observed in the treatment groups (Supplementary Fig. S7B). Survival rates following SNAP-TIL treatment were significantly higher (*p*=0.0002) than the control group (**Fig. 6B**). IHC of the lymphocyte population in the tumor periphery and core sections showed that SNAP-TIL significantly infiltrated the tumor (**Fig. 6C**, Supplementary Table S3) with a significantly higher number of CD8+ T cells than TIL REP. No significant difference was observed with CD4+ cells. A similar trend was observed by flow cytometry (**Fig. 6D**). Moreover, in the PDAC PDX model, tumor-bearing mice treated with SNAP-TIL showed significantly enhanced IFNγ expression in tumors compared to mice treated with TIL REP (**Fig. 6E**). The amount of IFNγ+ cells was significantly higher (3-fold) in tumors treated with SNAP-TIL compared to TIL REP-treated tumors. Apoptosis status measured by AnnexinV/7AAD staining revealed a higher number of early and late apoptotic/dead cells in SNAP-TIL-treated tumors and a lower number of live cells than in TIL REP-treated tumors (**Fig. 6F**). We checked the number of T cells in the spleen of mice by FACS and found an increase in the number of CD3+ CD8+ T cells compared to those in the TIL REP and control group (data not shown), indicating the proliferation of SNAP-TIL cells, possibly after the exposure to the neoantigens. Last, we measured neoantigen reactive T cell clones after SNAP-TIL treatment. CD45+ cells isolated from the PDAC PDX tumors were co-cultured with each neopeptide, and reactivity was measured using the IFNγ ELISpot assay. A total of 3 out of the 24 neopeptides (12.5%) produced a T cell response of sufficiently high magnitude compared to the baseline (**Fig. 6G**). Our *in vivo* data demonstrates SNAP-TIL to be reactive in both highly and poorly immunogenic cancers, with enhanced infiltration and antitumor activity.

**Figure 6.**
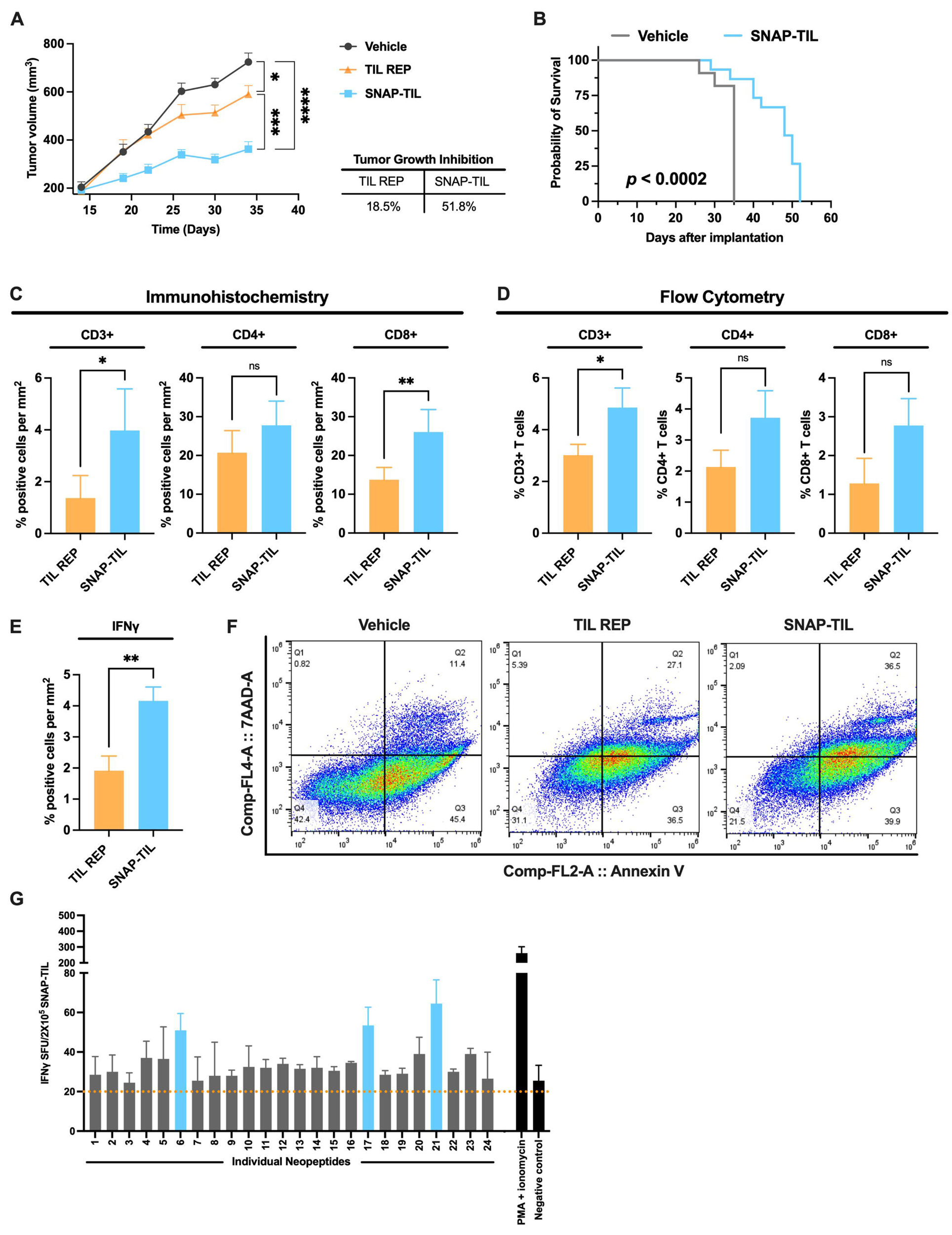
SNAP-TIL efficacy is superior to conventional TIL REP in the immunologically “cold” PDAC PDX model. (A) Female *hIL2*-NOG mice were subcutaneously implanted with propagated PDAC cells. Once tumors reached around 200 mm^3^, autologous SNAP-TIL, TIL REP, or vehicle was administered by retroorbital injection. Tumor volume was measured weekly and plotted as mean + SEM. Groups consisted of 12 - 14 mice. (B) Kaplan-Meier survival probability. (C) Quantification of CD3+, CD4+, and CD8+ cells in PDAC PDX by IHC. Each bar represents the mean ± SD, n = 5, \**p* < 0.05, \*\**p* < 0.01, ns, not significant, unpaired t-test. (D) Quantification of CD3+, CD4+, and CD8+ cells in PDAC PDX by flow cytometry. Each bar represents the mean ± SD, n = 3, \**p* < 0.05, ns, not significant, unpaired t-test. (E) Quantification of IFNγ positive cells in PDAC PDX by IHC. Each bar represents the mean ± SD, n = 3, \*\**p* < 0.01, unpaired t-test. (F) Annexin-V/7AAD surface expression of PDAC PDX by flow cytometry. Representative data from one of 3 experiments. (G) Detection of neoantigen reactive T cell clones after SNAP-TIL treatment in the PDAC PDX model. CD45+ cells isolated from the PDAC PDX tumors were co- cultured with each neopeptide, and reactivity was measured using the IFNγ ELISpot assay. Each bar represents the mean ± SD. Reactive neopeptides are indicated in blue. The dotted orange line represents the baseline.

## DISCUSSION

A critical element in successful TIL therapy is the recognition of tumor neoantigens; however, neoantigen-reactive T cells are often rare and exhausted in poorly immunogenic tumor types, complicating the development of highly reactive TIL products for patient treatment. Recently, Rosenberg et al. showed that neoantigenic stimulation enabled selective expansion of neoantigen-reactive TILs, not just by their frequencies but also by broadening their clonal repertoires, overall improving the reactivity, phenotypes, and functions of TILs (8). Consequently, neoantigen selection is a key step in developing TIL therapy products. In this study, we introduce our new neoantigen identification technology platform - SNAP, which improves neoantigen prediction to generate autologous TILs enriched for T cells that specifically recognize clonal neoantigenic epitopes. The SNAP-TIL produced post-REP across all tumor types was a near-homogenous mixture of CD4 and CD8 cells, with an approximate 3:1 ratio of effector: central memory cells. While most cells in the SNAP-TIL product were in a differentiated state, there was a small reservoir of TSMCs known for their ability to self-renew, expand robustly, and exhibit strong antitumor activity *in vitro* and *in vivo* (31). SNAP-TIL exhibited greater efficacy in immune infiltration than nonspecific TIL REP only using *ex vivo* tumoroids. It also proved highly selective for autologous tumoroids with no cross-reactivity to allogenic tumoroids or NAT, as observed with TIL REP only. SNAP-TIL also displayed superior activation with robust immunogenicity and on-target tumor cell killing in highly immunogenic and poorly immunogenic tumors, with 70% and 50% TGI in melanoma and PDAC PDX models, respectively. These results demonstrate that TIL infusion products enriched with neoantigen reactivity before *in vitro* expansion and infusion improve the antitumor response rate over TIL REP only.

Our study raises several important points. Neoantigen discovery relying solely on WES DNA analysis and/or *in silico* prediction may miss critically important neoantigenic epitopes, particularly in the case of immunologically cold tumors. Indeed, neoantigens can arise from any genomic variation, differential expression, and post-translational modification (32–43). A recent study reported that neoantigens inferred from RNA analysis might represent a relevant target for ACT (44). Integrating WES DNA analysis with mRNA expression data will improve the likelihood of identifying immunogenic neoantigens, potentially improving immunotherapeutic approaches targeting neoantigens.

Another important point of consideration is the time needed to generate SNAP-TIL and its potential impact on cancer patients. T-cell expansion for infusion is time-consuming, taking weeks to prepare, and time is limited for many cancer patients. Unlike the TIL bulk preparation, the SNAP platform requires additional steps, translating to time increments to release the final TIL product. High-efficiency SNAP-TIL production requires 6 to 8 weeks, which may not be available to late-stage cancer patients. We are improving our protocol to reduce the preparation time by accelerating the sequencing, analysis, and binding assay steps in our protocol. In parallel, we are using the same approach to develop individualized peptide-based or mRNA vaccines that stimulate neoantigen-specific T cells. Vaccine preparation time is likely more amenable to late-stage patients as the education and REP steps for SNAP-TIL would be unnecessary.

Lastly, our study was not powered to investigate the effect of SNAP-TIL combined with immune checkpoint inhibitors (ICIs). TIL therapy combined with programmed cell death ligand-1 (PD-L1) has been shown to prolong survival in patients with metastatic disease (45), and some have suggested that combining TIL therapy with PD-1 may increase TIL fold expansion after infusion across multiple solid cancers (45,46). Investigating the efficacy of SNAP-TIL in combination with ICIs is a priority for the immediate future.

This study demonstrates the feasibility of our novel ‘Personalized Neoantigen Pipeline’ approach to developing a high-efficiency product in ACT or other immunotherapies reliant on neoantigen targeting. While this was a low-powered study based on a relatively small cohort of patient samples, the SNAP-TIL platform appears safe and feasible, generating a durable antitumor immune response even for immunosuppressive cancers. These early results warrant further development of SNAP-TIL, alone or combined with other therapies to treat solid cancers.

## Acknowledgments

This work was supported by the Dorrance Family Research Fund. The content and views expressed in this article are solely those of the authors and do not necessarily represent the official policy or position of the funding sponsors. The National Institutes of Health MHC Tetramer Core Facility (Emory University, Atlanta, USA) kindly provided the MHC II reagents used in this study.

## Authors’ Contributions

**T. Halder:** Data curation, formal analysis, investigation, methodology, validation, visualization, writing – review and editing. **E. Kelly:** Resources, software. **J. Soria-Bustos:** Resources. **T Bargenquast:** Resources, writing – review and editing. **R. Rodriquez del Villar:** Resources, formal analysis. **A. Weston:** Writing – review and editing. **T. Thode:** Writing – review and editing. **S. Ng:** Writing – review and editing. **M. Kaadige:** Formal analysis, visualization, writing – review and editing. **E. Borazanci:** Resources. **M. Gordon:** Resources. **J. Moser:** Resources. **F. Tsai:** Resources. **S. Priceman:** Writing – review & editing. **S. Forman:** Writing – review & editing. **J. Altin:** Resources, software. **R. Soldi:** Conceptualization, data curation, formal analysis, investigation, methodology, project administration, resources, software, supervision, validation, visualization, writing – original draft, writing – review and editing. **S. Sharma:** Conceptualization, formal analysis, funding acquisition, investigation, project administration, resources, supervision, validation, writing – review and editing.

## Data availability statement

The data supporting this study’s findings are available from the corresponding author, RS, upon reasonable request.

