## Supplementary figures for "Preclinical Proof of Concept for a personalized SNAP^TM^-TIL (Specific Neo-Antigen Peptides -TIL) therapy platform"

**Supplementary Figure S1**

Immunophenotyping of SNAP-TIL and TIL REP products post-REP. Dotplots of flow cytometry data are demonstrating the strategy to identify different immune cell subtypes: (A) CD3+ cells, (B) CD4+ and CD8+ cells, (C) NK cells, (D) B cells, (E) activated T cells, (F) effector and central memory T cells, (G) Treg cells, or (H) Epcam+ cells. The data shows the characterization of SNAPTILs generated from the representative (1) melanoma, (2, 4) PDAC, (3) uveal melanoma, (5) CRC patients.

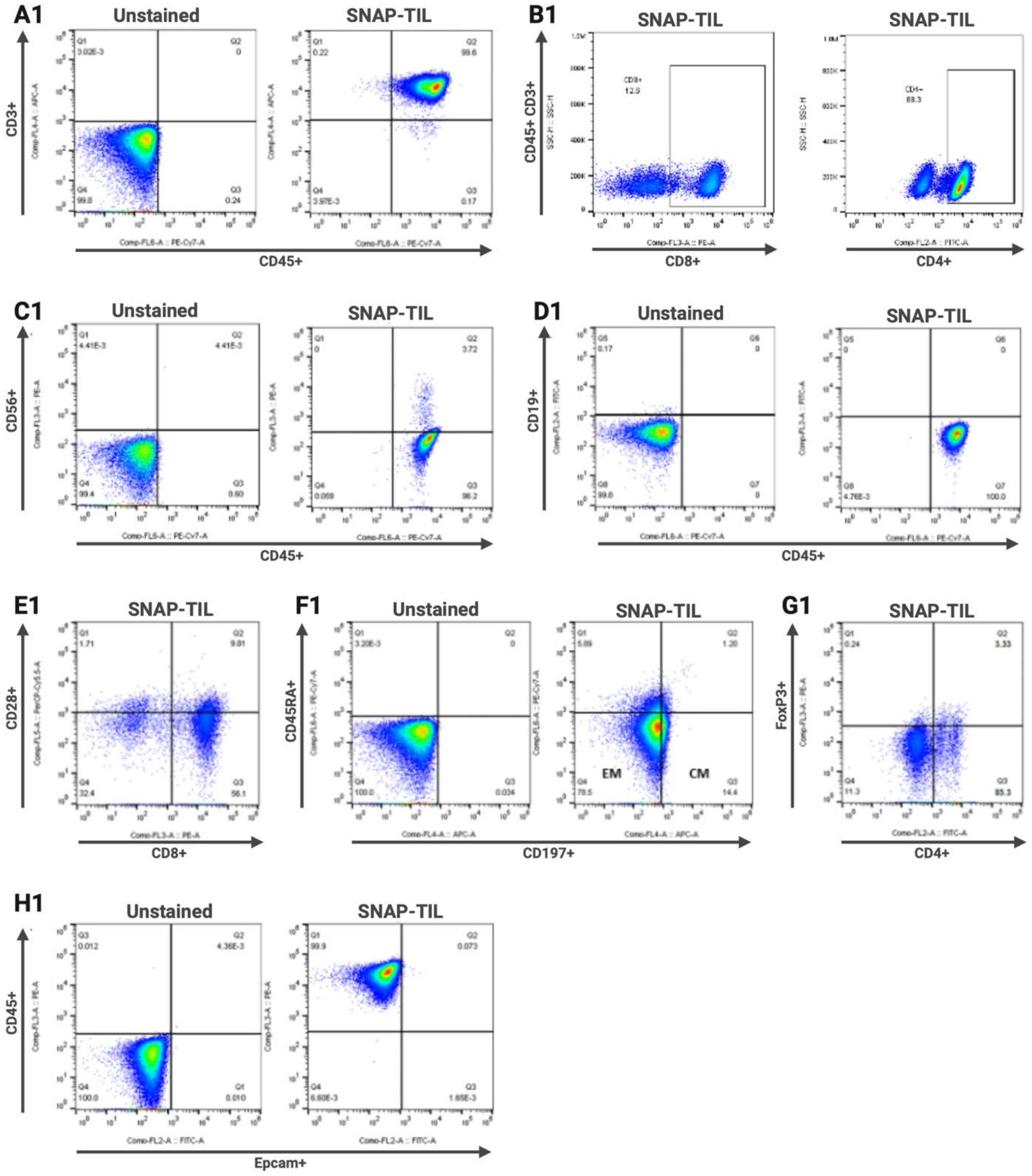

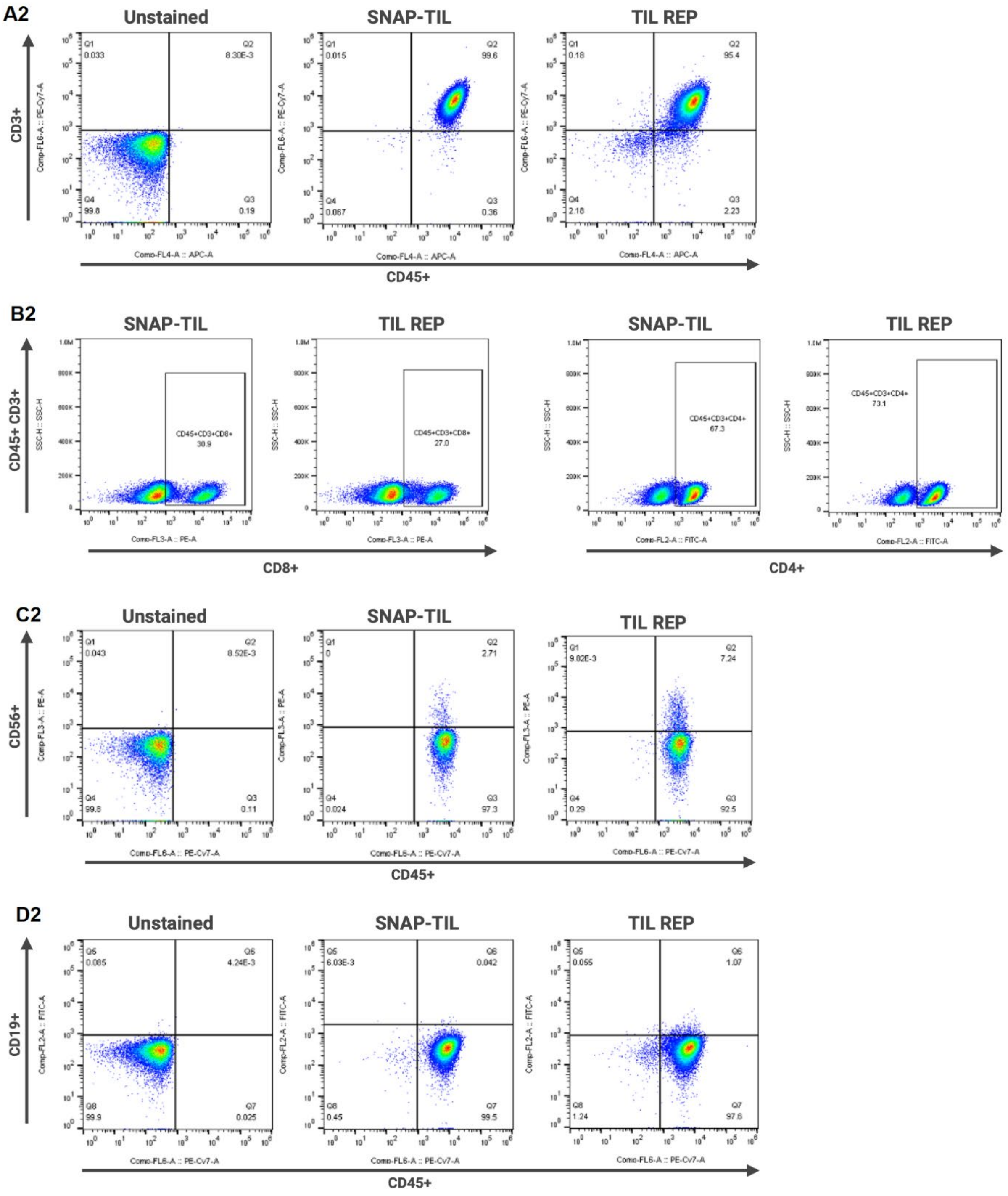
A2
Unstained
SNAP-TIL
TIL REP
CD3+
Comp-FL6-A :: PE-Cy7-A
Q1 0.033
Q2 8.30E-3
Q4 99.8
Q3 0.19
Q1 0.015
Q2 99.6
Q4 0.067
Q3 0.36
Q1 0.18
Q2 95.4
Q4 2.18
Q3 2.23
Comp-FL4-A :: APC-A
CD45+
B2
SNAP-TIL
TIL REP
SNAP-TIL
TIL REP
CD45+ CD3+
SSC-H :: SSC-H
CD45+CD3+CD8+
30.9
CD45+CD3+CD8+
27.0
CD45+CD3+CD4+
67.3
CD45+CD3+CD4+
73.1
Comp-FL3-A :: PE-A
Comp-FL2-A :: FITC-A
CD8+
CD4+
C2
Unstained
SNAP-TIL
TIL REP
CD56+
Comp-FL3-A :: PE-A
Q1 0.043
Q2 8.52E-3
Q4 99.8
Q3 0.11
Q1 0
Q2 2.71
Q4 0.024
Q3 97.3
Q1 9.82E-3
Q2 7.24
Q4 0.29
Q3 92.5
Comp-FL6-A :: PE-Cy7-A
CD45+
D2
Unstained
SNAP-TIL
TIL REP
CD19+
Comp-FL2-A :: FITC-A
Q5 0.085
Q6 4.24E-3
Q8 99.9
Q7 0.025
Q5 6.03E-3
Q6 0.042
Q8 0.45
Q7 99.5
Q5 0.055
Q6 1.07
Q8 1.24
Q7 97.6
Comp-FL6-A :: PE-Cy7-A
CD45+

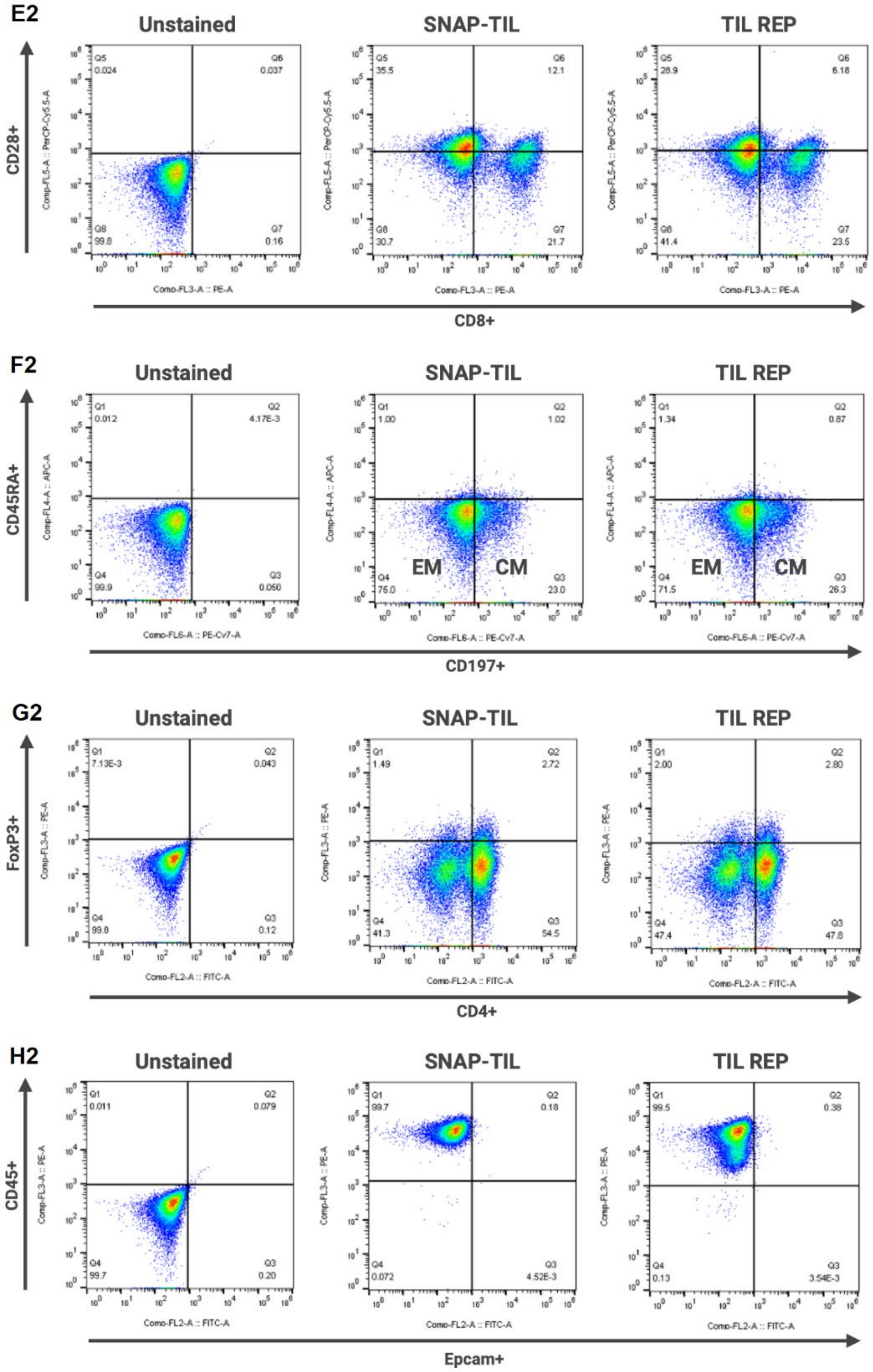

E2
Unstained
SNAP-TIL
TIL REP
CD28+
CD8+
Q5 0.024
Q6 0.037
Q8 99.8
Q7 0.16
Q5 35.5
Q6 12.1
Q8 30.7
Q7 21.7
Q5 28.9
Q6 6.18
Q8 41.4
Q7 23.5
F2
Unstained
SNAP-TIL
TIL REP
CD45RA+
CD197+
Q1 0.012
Q2 4.17E-3
Q4 99.9
Q3 0.050
Q1 1.00
Q2 1.02
Q4 75.0
Q3 23.0
Q1 1.34
Q2 0.87
Q4 71.5
Q3 26.3
EM
CM
G2
Unstained
SNAP-TIL
TIL REP
FoxP3+
CD4+
Q1 7.13E-3
Q2 0.043
Q4 99.8
Q3 0.12
Q1 1.49
Q2 2.72
Q4 41.3
Q3 54.5
Q1 2.00
Q2 2.80
Q4 47.4
Q3 47.8
H2
Unstained
SNAP-TIL
TIL REP
CD45+
Epcam+
Q1 0.011
Q2 0.079
Q4 99.7
Q3 0.20
Q1 99.7
Q2 0.18
Q4 0.072
Q3 4.52E-3
Q1 99.5
Q2 0.38
Q4 0.13
Q3 3.54E-3

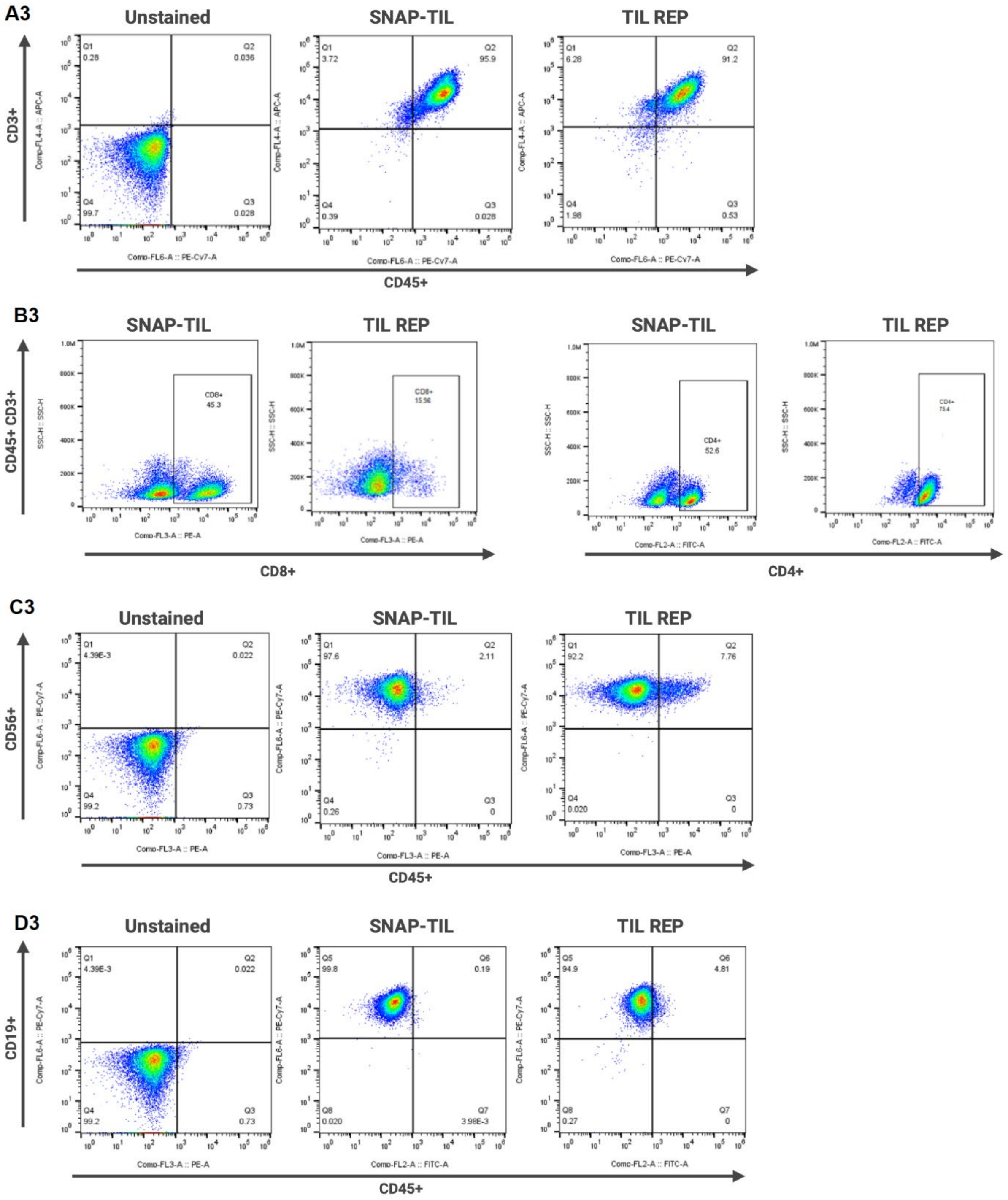
A3
Unstained
SNAP-TIL
TIL REP
CD3+
Comp-FL4-A :: APC-A
Q1 0.28
Q2 0.036
Q4 99.7
Q3 0.028
Q1 3.72
Q2 95.9
Q4 0.39
Q3 0.028
Q1 6.28
Q2 91.2
Q4 1.98
Q3 0.53
Comp-FL6-A :: PE-Cy7-A
CD45+
B3
SNAP-TIL
TIL REP
SNAP-TIL
TIL REP
CD45+ CD3+
SSC-H :: SSC-H
CD8+ 45.3
CD8+ 15.96
CD4+ 52.6
Comp-FL3-A :: PE-A
Comp-FL2-A :: FITC-A
CD8+
CD4+
C3
Unstained
SNAP-TIL
TIL REP
CD56+
Comp-FL6-A :: PE-Cy7-A
Q1 4.39E-3
Q2 0.022
Q4 99.2
Q3 0.73
Q1 97.6
Q2 2.11
Q4 0.26
Q3 0
Q1 92.2
Q2 7.76
Q4 0.020
Q3 0
Comp-FL3-A :: PE-A
CD45+
D3
Unstained
SNAP-TIL
TIL REP
CD19+
Comp-FL6-A :: PE-Cy7-A
Q1 4.39E-3
Q2 0.022
Q4 99.2
Q3 0.73
Q5 99.8
Q6 0.19
Q8 0.020
Q7 3.98E-3
Q5 94.9
Q6 4.81
Q8 0.27
Q7 0
Comp-FL3-A :: PE-A
Comp-FL2-A :: FITC-A
CD45+

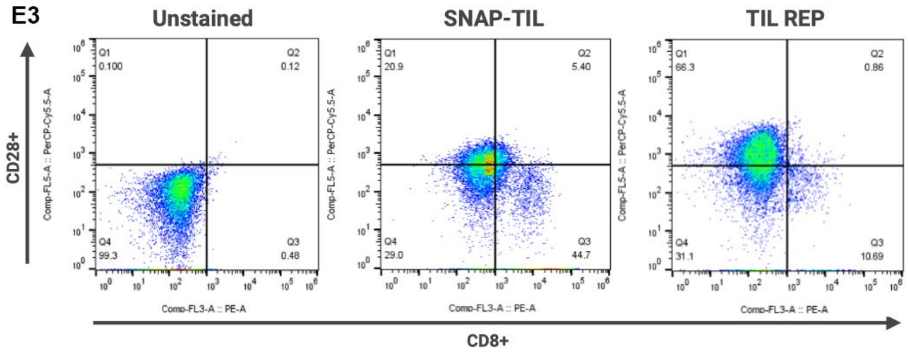
E3
Unstained
SNAP-TIL
TIL REP
CD28+
CD8+
Comp-FL5-A :: PerCP-Cy5.5-A
Comp-FL3-A :: PE-A
Q1
0.100
Q2
0.12
Q4
99.3
Q3
0.48
Q1
20.9
Q2
5.40
Q4
29.0
Q3
44.7
Q1
66.3
Q2
0.86
Q4
31.1
Q3
10.69

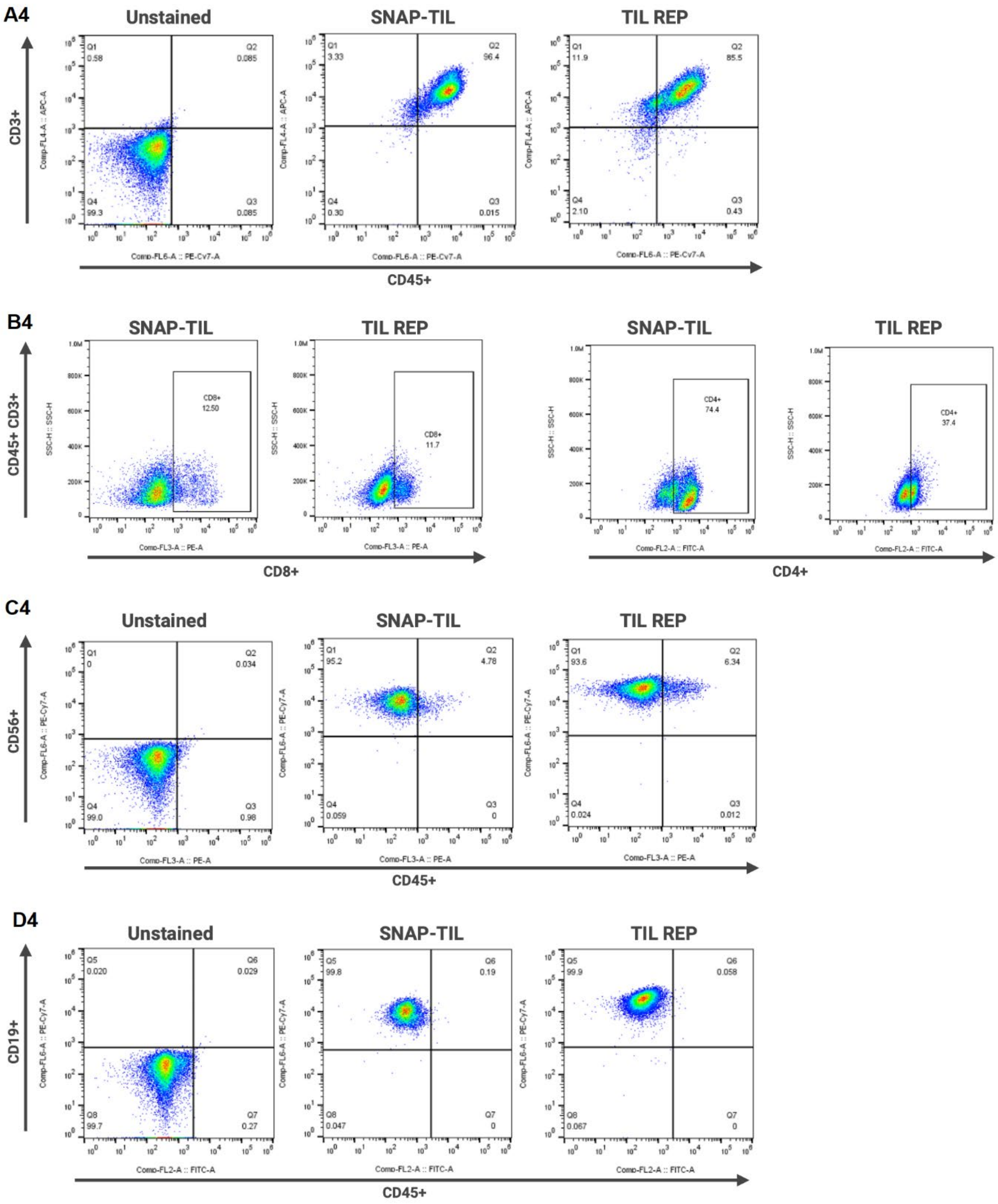

A4
Unstained
SNAP-TIL
TIL REP
CD3+
Comp-FL4-A :: APC-A
Q1
0.58
Q2
0.085
Q4
99.3
Q3
0.085
Q1
3.33
Q2
96.4
Q4
0.30
Q3
0.015
Q1
11.9
Q2
85.5
Q4
2.10
Q3
0.43
Comp-FL6-A :: PE-Cy7-A
CD45+
B4
SNAP-TIL
TIL REP
SNAP-TIL
TIL REP
CD45+ CD3+
SSC-H :: SSC-H
CD8+
12.50
CD8+
11.7
CD4+
74.4
CD4+
37.4
Comp-FL3-A :: PE-A
Comp-FL2-A :: FITC-A
CD8+
CD4+
C4
Unstained
SNAP-TIL
TIL REP
CD56+
Comp-FL6-A :: PE-Cy7-A
Q1
0
Q2
0.034
Q4
99.0
Q3
0.98
Q1
95.2
Q2
4.78
Q4
0.059
Q3
0
Q1
93.6
Q2
6.34
Q4
0.024
Q3
0.012
Comp-FL3-A :: PE-A
CD45+
D4
Unstained
SNAP-TIL
TIL REP
CD19+
Comp-FL6-A :: PE-Cy7-A
Q5
0.020
Q6
0.029
Q8
99.7
Q7
0.27
Q5
99.8
Q6
0.19
Q8
0.047
Q7
0
Q5
99.9
Q6
0.058
Q8
0.067
Q7
0
Comp-FL2-A :: FITC-A
CD45+

E4

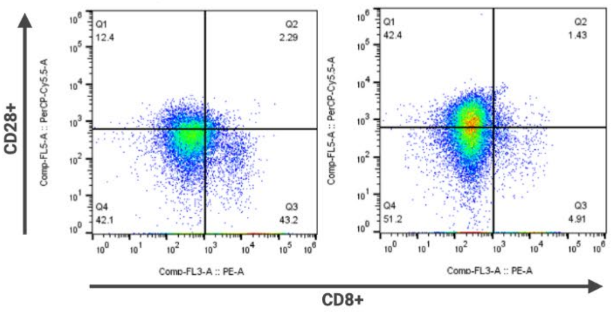
SNAP-TIL
TIL REP
CD28+
CD8+
Comp-FL5-A :: PerCP-Cy5.5-A
Comp-FL3-A :: PE-A
Q1
12.4
Q2
2.29
Q4
42.1
Q3
43.2
Q1
42.4
Q2
1.43
Q4
51.2
Q3
4.91

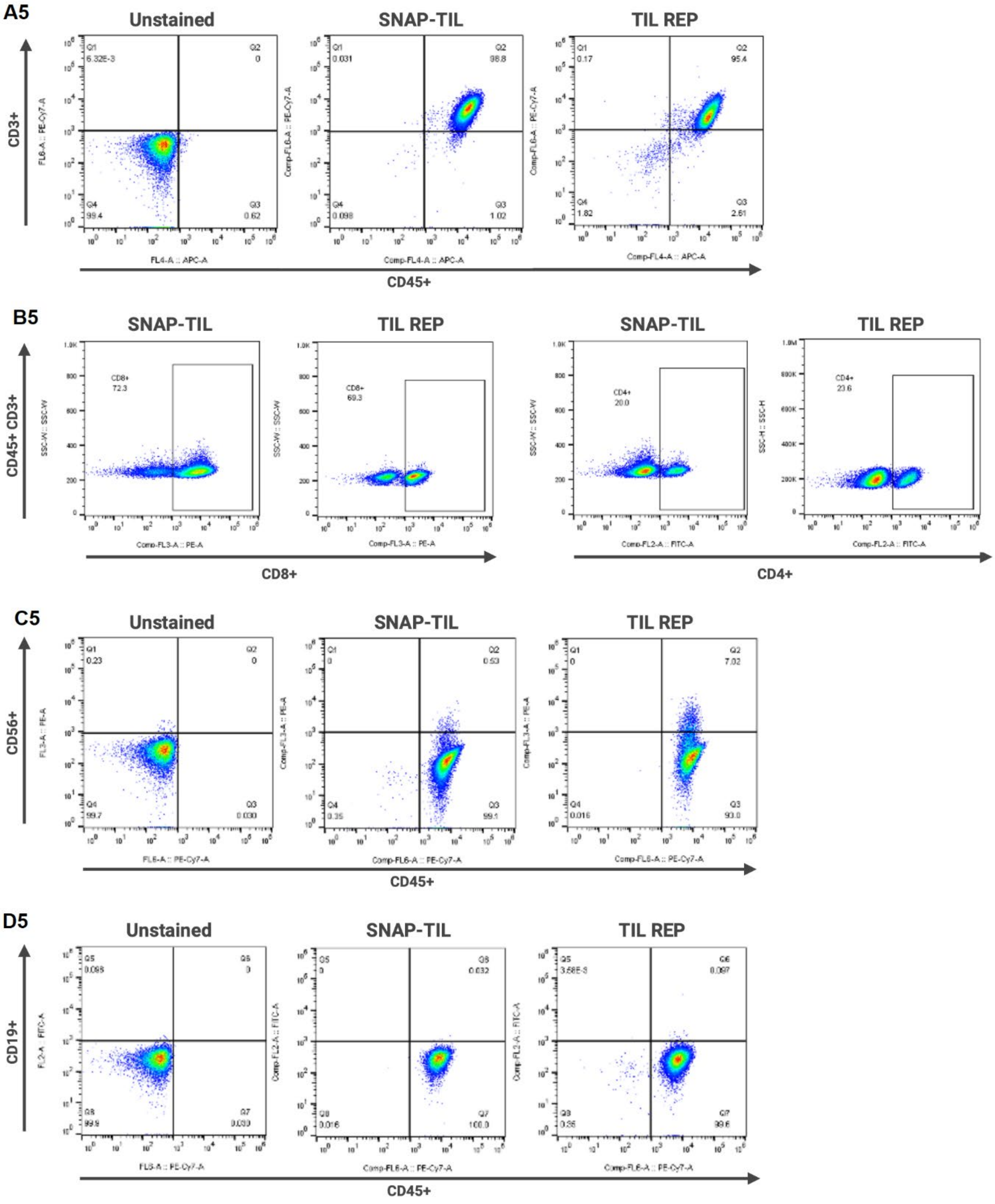
A5
Unstained
SNAP-TIL
TIL REP
CD3+
CD45+
B5
CD45+ CD3+
CD8+
CD4+
C5
CD56+
D5
CD19+

E5

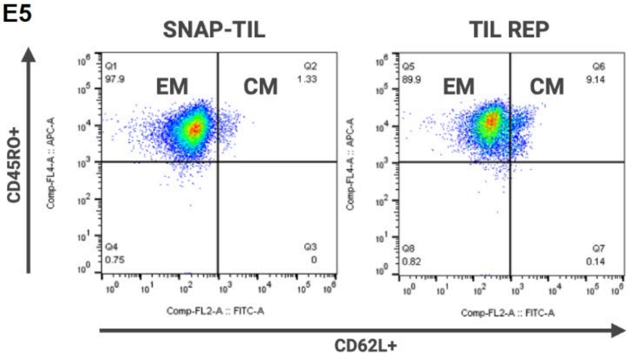
SNAP-TIL
TIL REP
EM
CM
EM
CM
Q1
97.9
Q2
1.33
Q4
0.75
Q3
0
Q5
89.9
Q6
9.14
Q8
0.82
Q7
0.14
Comp-FL4-A :: APC-A
Comp-FL2-A :: FITC-A
CD45RO+
CD62L+

F5

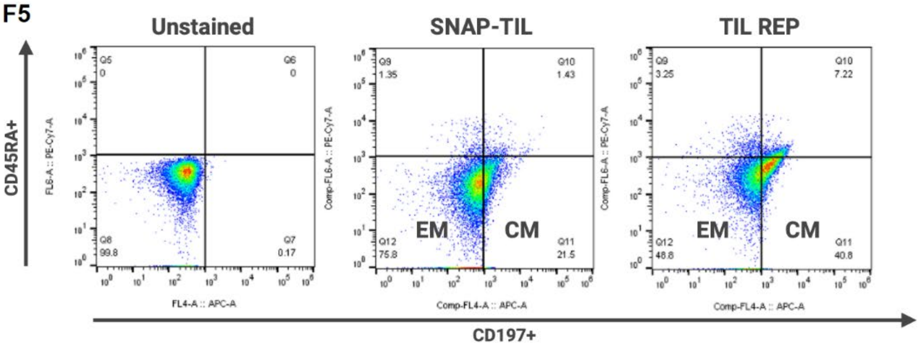
Unstained
SNAP-TIL
TIL REP
Q5
0
Q6
0
Q8
99.8
Q7
0.17
Q9
1.35
Q10
1.43
Q12
75.8
Q11
21.5
Q9
3.25
Q10
7.22
Q12
48.8
Q11
40.8
EM
CM
EM
CM
FL6-A :: PE-Cy7-A
Comp-FL6-A :: PE-Cy7-A
FL4-A :: APC-A
Comp-FL4-A :: APC-A
CD45RA+
CD197+

G5

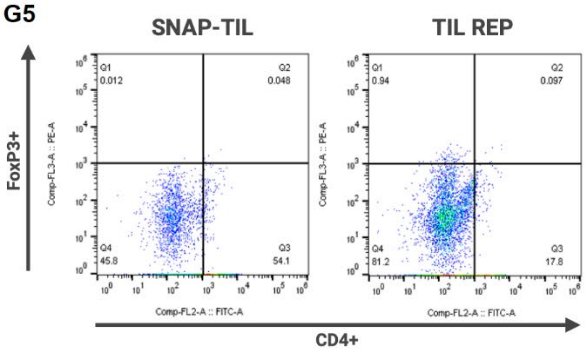
SNAP-TIL
TIL REP
Q1
0.012
Q2
0.048
Q4
45.8
Q3
54.1
Q1
0.94
Q2
0.097
Q4
81.2
Q3
17.8
Comp-FL3-A :: PE-A
Comp-FL2-A :: FITC-A
FoxP3+
CD4+

H5

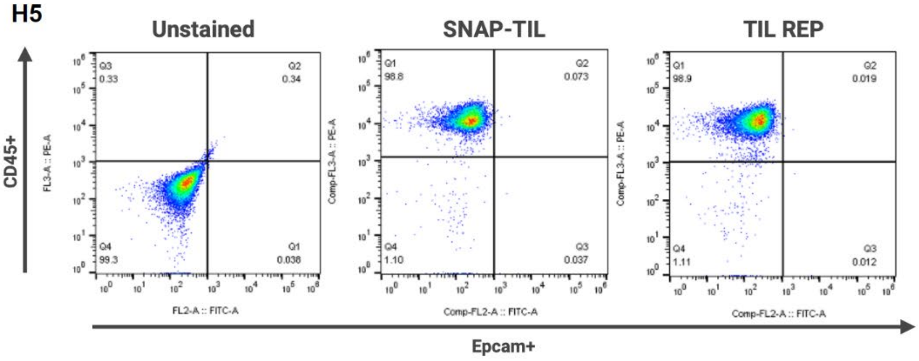
Unstained
SNAP-TIL
TIL REP
Q3
0.33
Q2
0.34
Q4
99.3
Q1
0.036
Q1
98.8
Q2
0.073
Q4
1.10
Q3
0.037
Q1
98.9
Q2
0.019
Q4
1.11
Q3
0.012
FL3-A :: PE-A
Comp-FL3-A :: PE-A
FL2-A :: FITC-A
Comp-FL2-A :: FITC-A
CD45+
Epcam+

**Supplementary Figure S2**

Representative T cell UMAP sub-clustering analysis in PDAC patient's derived SNAP-TIL. Classification of sub-clusters: (A) CD45+ and CD3+ cells, (B) CD8+ cells, (C) CD4+ cells, (D) NK cells, (E) exhaustion T cell markers, (F) cytotoxic T cell markers, and (G) proliferating T cell markers. Each dot refers to a cell, and the scale is marker expression.

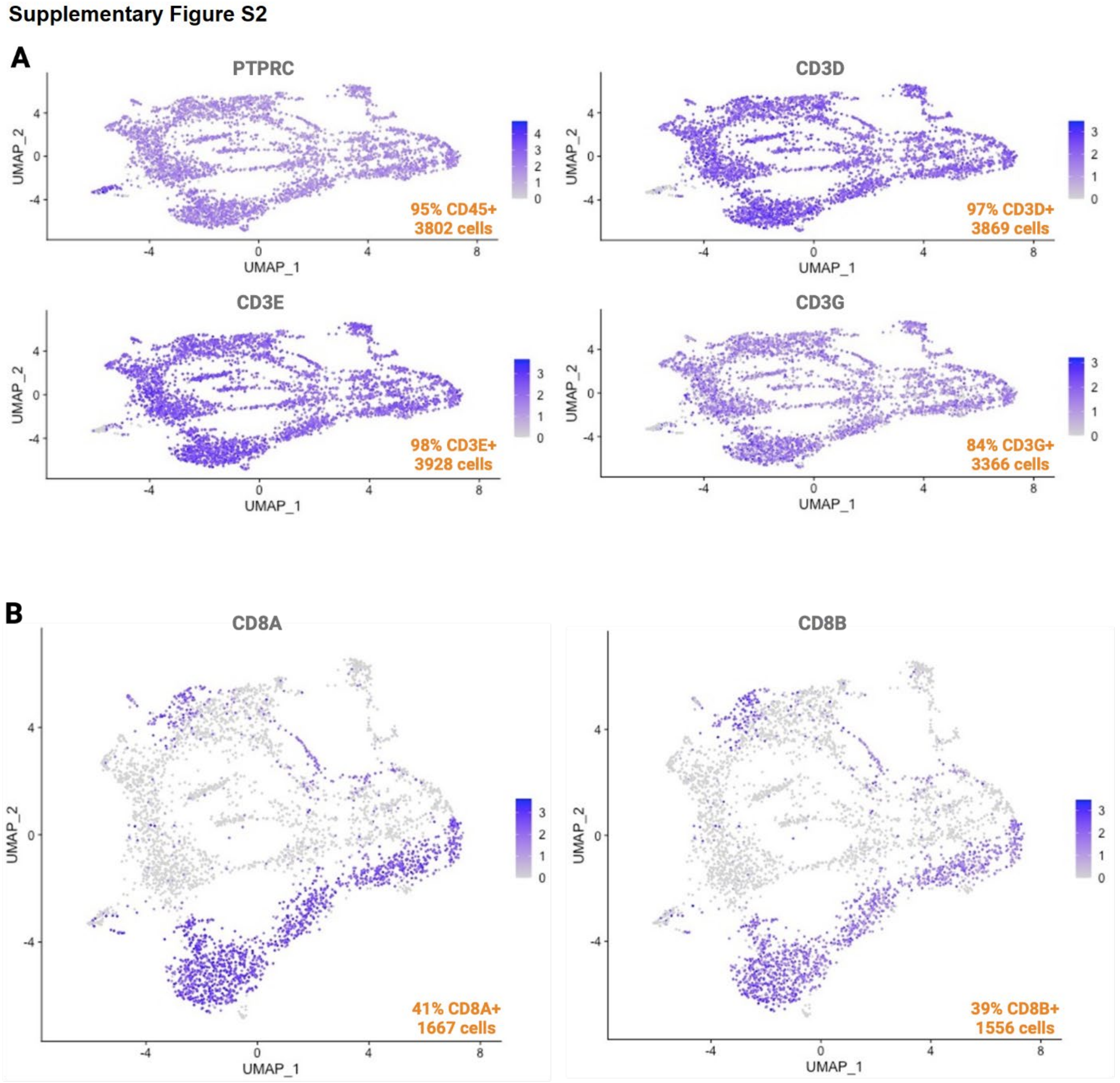

**C**

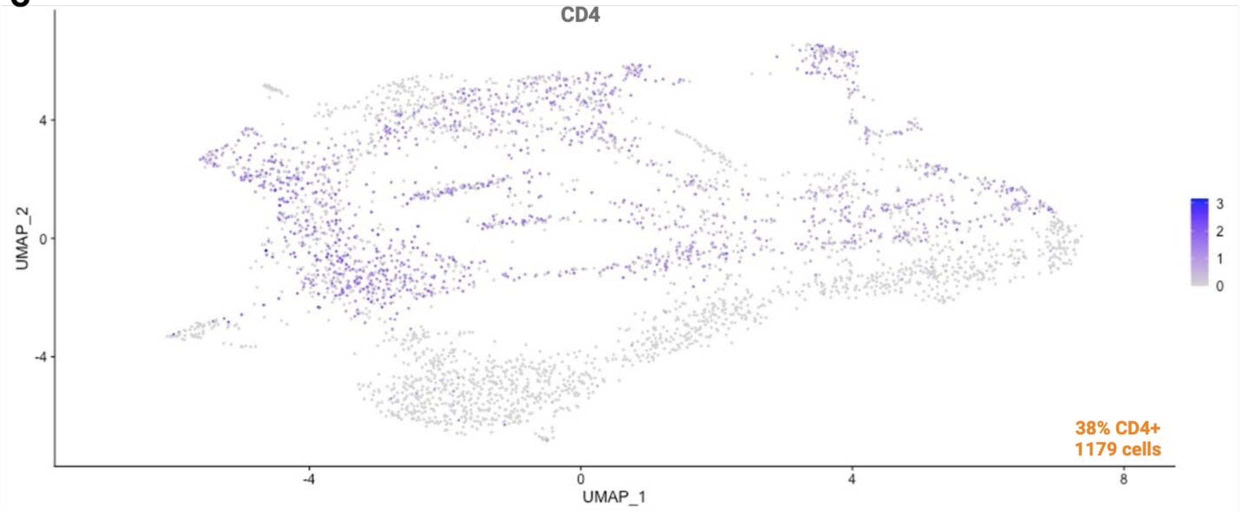
CD4
UMAP_2
UMAP_1
38% CD4+
1179 cells

**D**

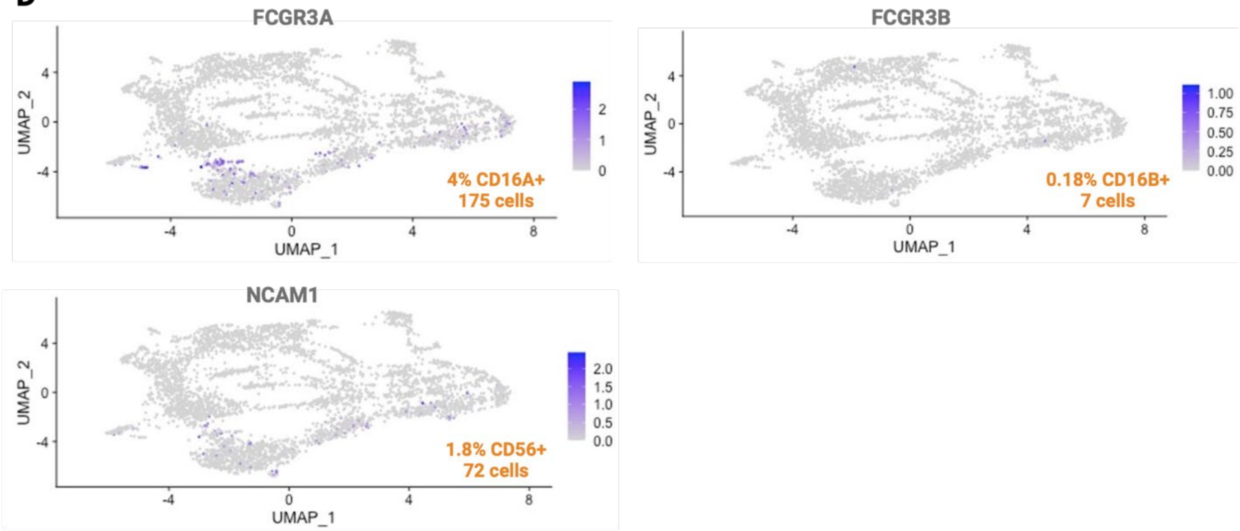
FCGR3A
4% CD16A+
175 cells
FCGR3B
0.18% CD16B+
7 cells
NCAM1
1.8% CD56+
72 cells
UMAP_2
UMAP_1

E

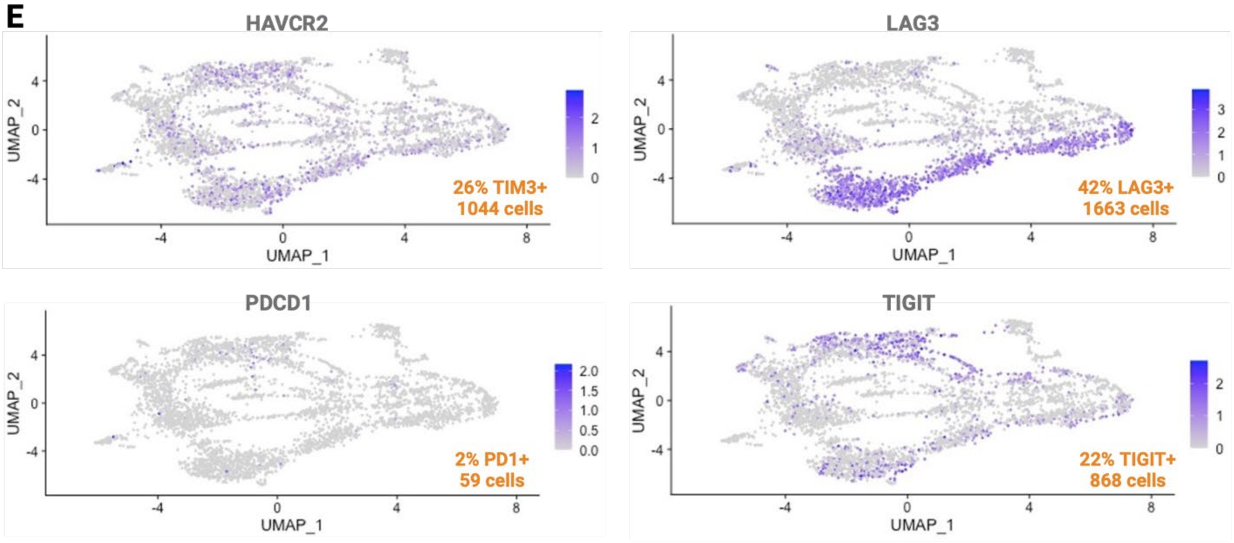
HAVCR2
LAG3
PDCD1
TIGIT
UMAP_2
UMAP_1
26% TIM3+
1044 cells
42% LAG3+
1663 cells
2% PD1+
59 cells
22% TIGIT+
868 cells

F

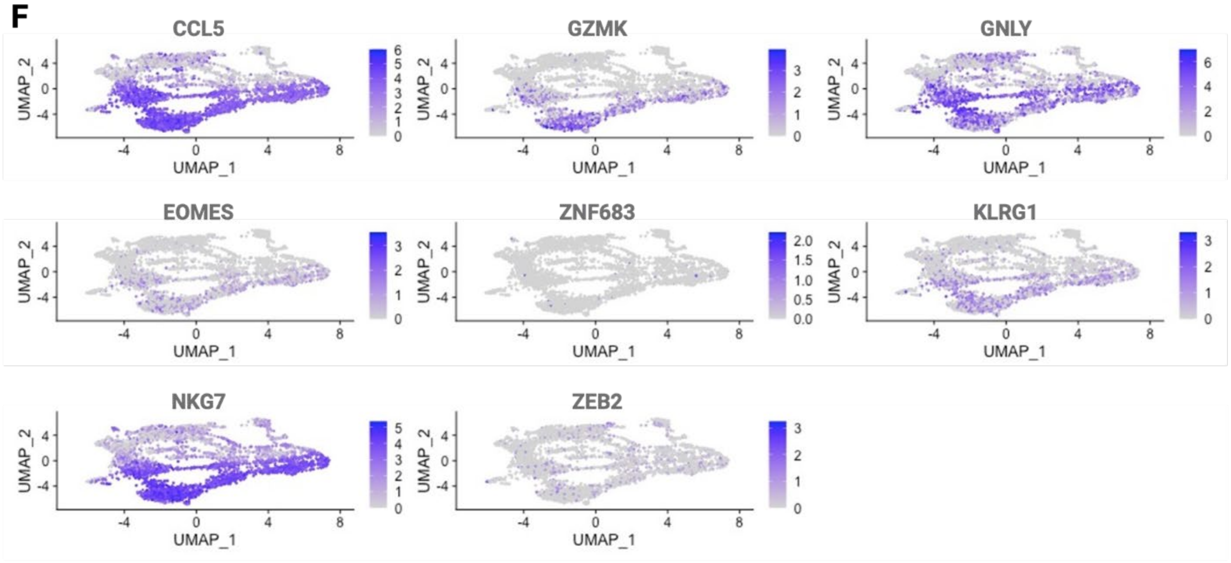
CCL5
GZMK
GNLY
EOMES
ZNF683
KLRG1
NKG7
ZEB2
UMAP_2
UMAP_1

G

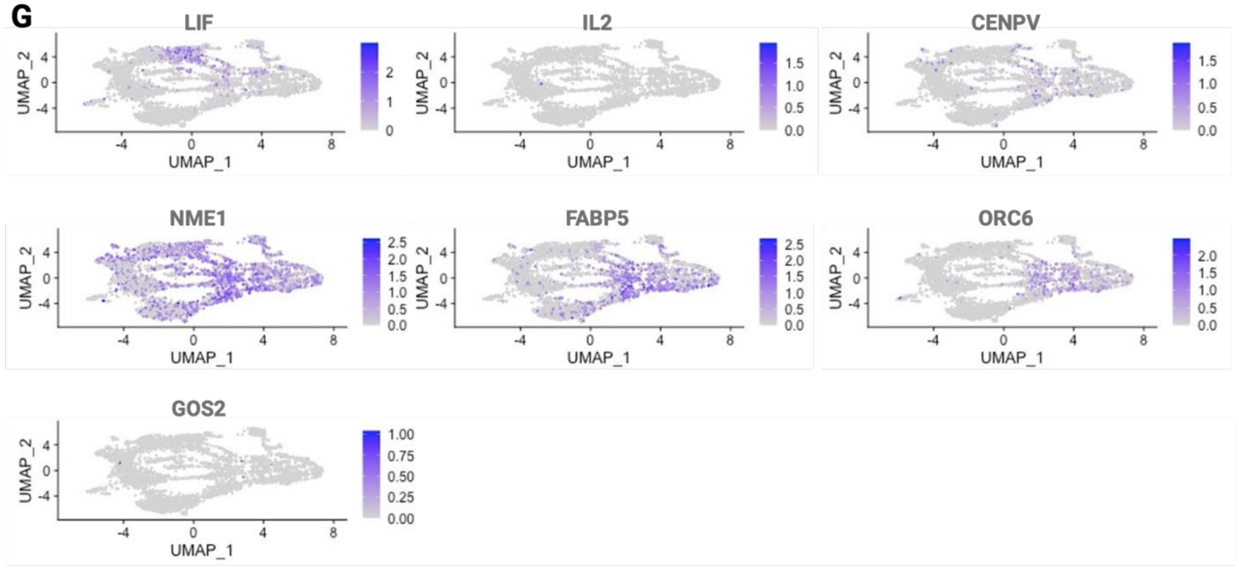
LIF
IL2
CENPV
NME1
FABP5
ORC6
GOS2
UMAP_2
UMAP_1

**Supplementary Figure S3.**

Comparison of immune reactivity of SNAP-TIL expanded with PepSeq or *in silico* peptide pools. **(A)** Stimulation of SNAP-TIL in the presence of individual neopeptides for CRC, uveal melanoma, and PDAC measured by IFNγ ELISpot assay. Bars represent mean ± SD, n = 2 replicates. (B) MSD measure of IFNγ levels in SNAP-TIL expanded with PepSeq or *in silico* peptide pool. Each dot represents one peptide; the horizontal bar indicates mean values, n = 7 – 19, *, $p < 0.05$, unpaired t-test. (C) Statistical estimation plot of the IFNγ level mean values from PepSeq and *in silico* peptides. (D) IFNγ level fold change in SNAP-TIL expanded by PepSeq pool, *in silico* pool, or a combined pool for CRC, melanoma, or PDAC. In the box plot diagram, the median is represented with a line, the interquartile with a box, and the minimum and maximum of the data with the whiskers, n = x, *, $p < 0.05$, ****, $p < 0.0001$, one-way ANOVA with Dunnett's post-hoc test.

**Supplementary Figure S3**

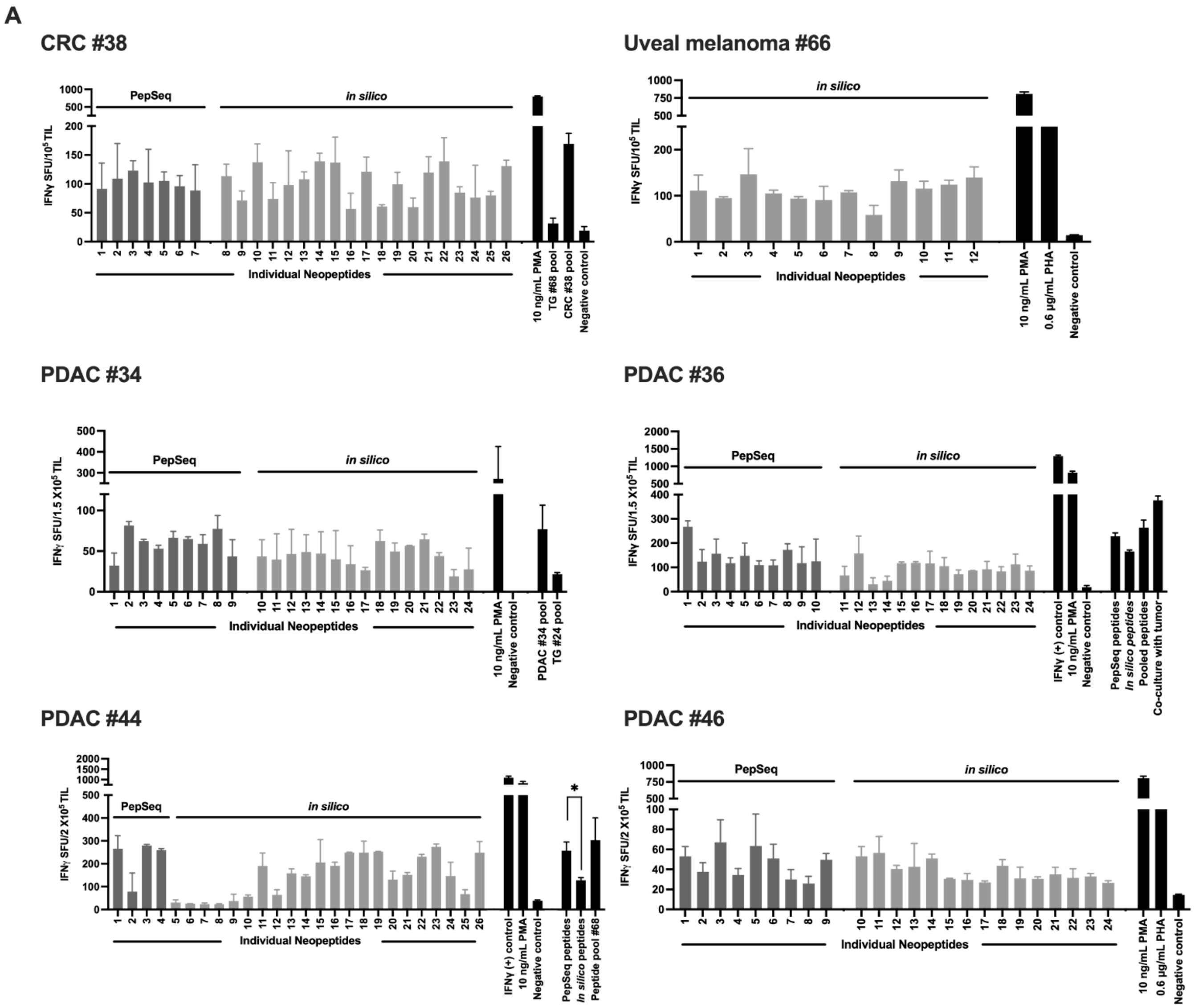

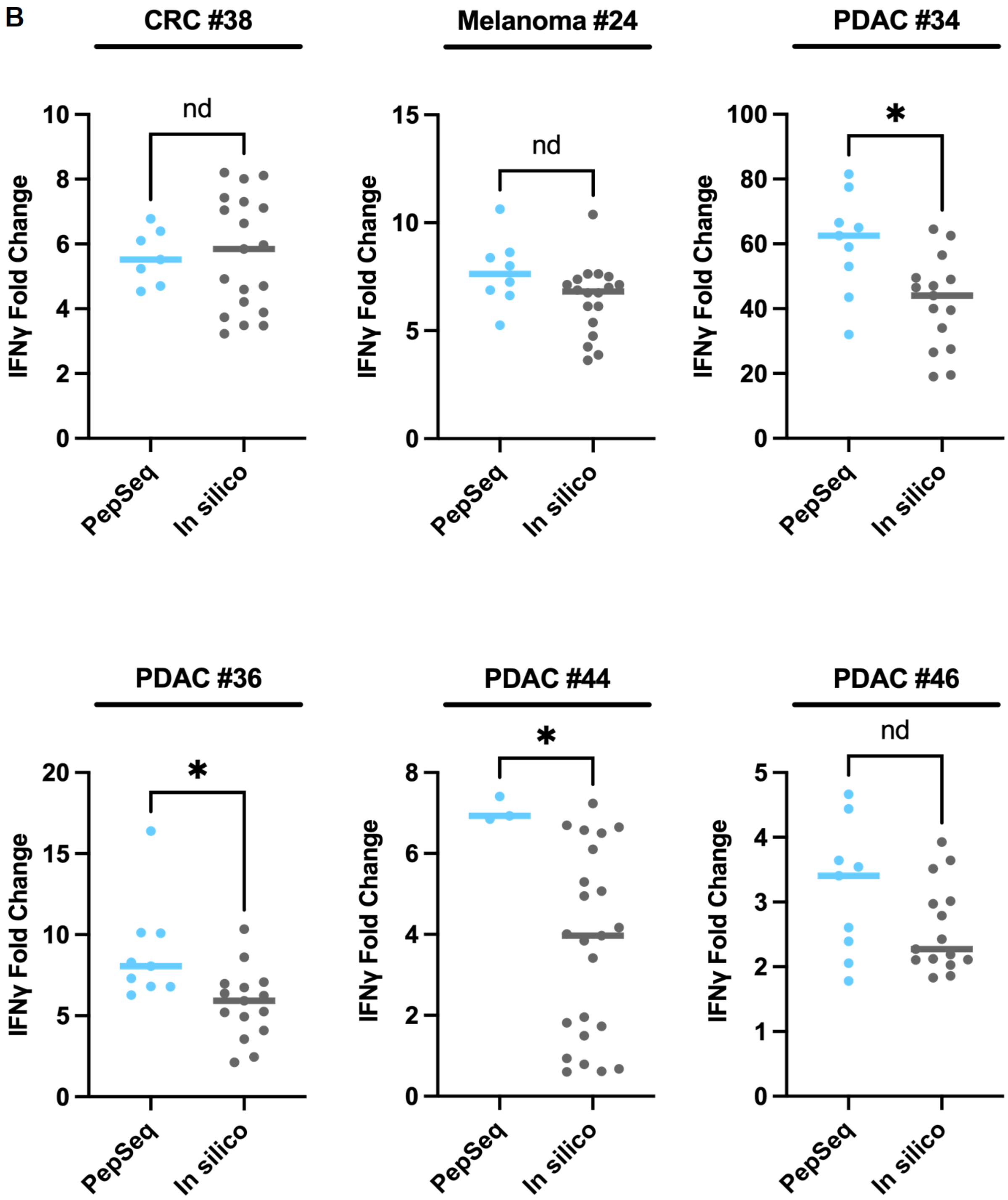
B
CRC #38
Melanoma #24
PDAC #34
PDAC #36
PDAC #44
PDAC #46
IFNγ Fold Change
nd
*
PepSeq
In silico
0
2
4
6
8
10
5
15
20
40
60
80
100
1
3

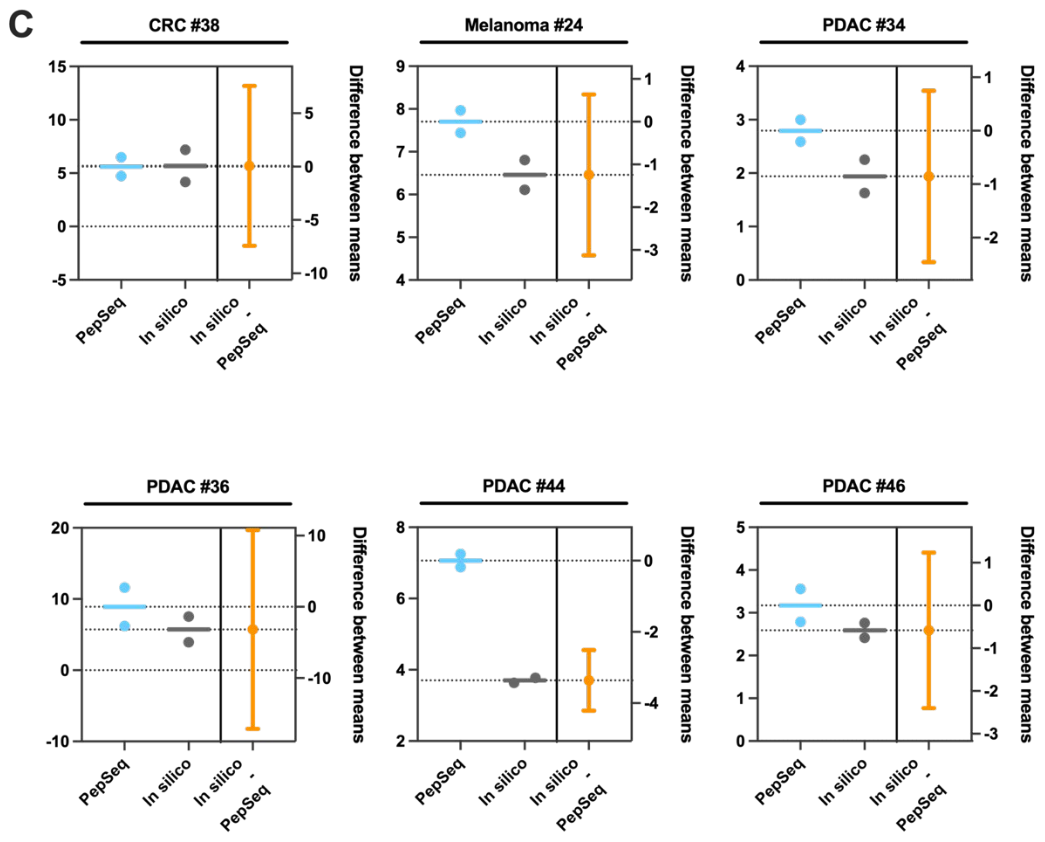
C
CRC #38
Melanoma #24
PDAC #34
PDAC #36
PDAC #44
PDAC #46
Difference between means
PepSeq
In silico
In silico - PepSeq

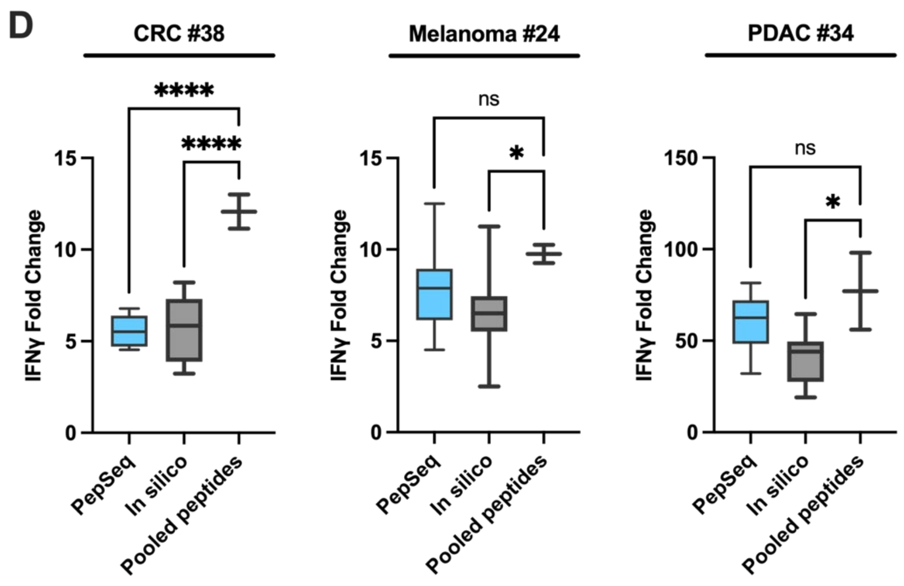
D
CRC #38
Melanoma #24
PDAC #34
****
****
ns
*
IFNγ Fold Change
PepSeq
In silico
Pooled peptides

**Supplementary Figure S4.**

TIL infiltration into melanoma or PDAC tumoroids after ‘educated’ with neopeptides (SNAP-TIL) outperforms nonspecific, conventional stimulation with TIL REP. Bar graphs represent mean ± SD, n = 3 - 5, *$p < 0.05$, **$p < 0.01$, ***$p < 0.001$, ****$p < 0.0001$, ns, not significant, ordinary one-way ANOVA with Tukey’s post-hoc test.

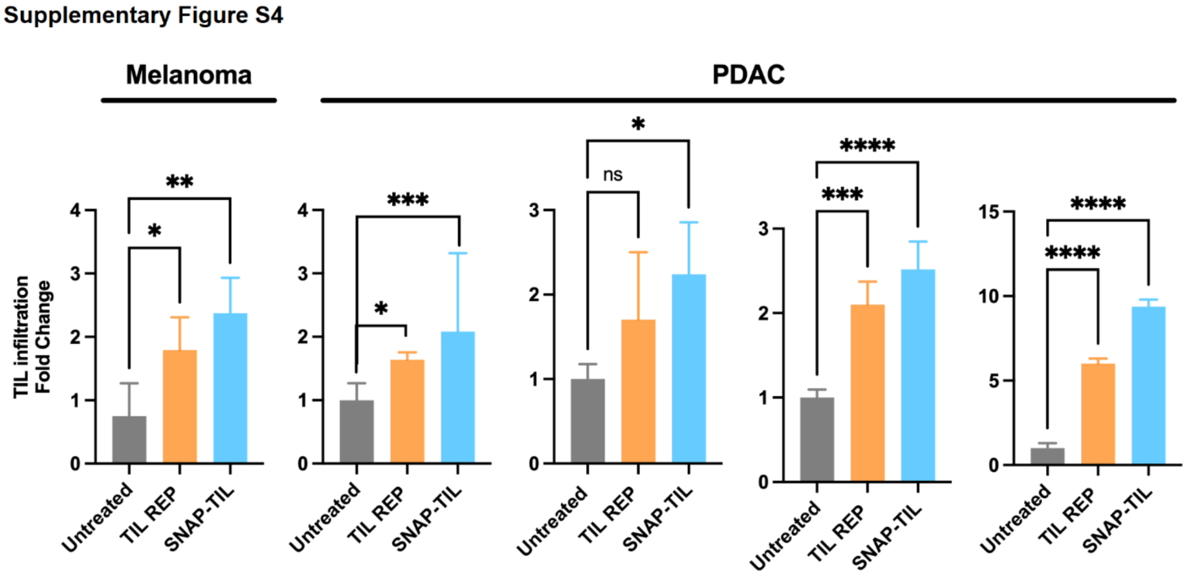

**Supplementary Figure S5**

IFNγ and granzyme B expression fold change in TILs co-cultured with autologous *ex-vivo* tumoroids: (A) CRC, (B) melanoma, (C) NSCLC, and (D-G) PDAC.

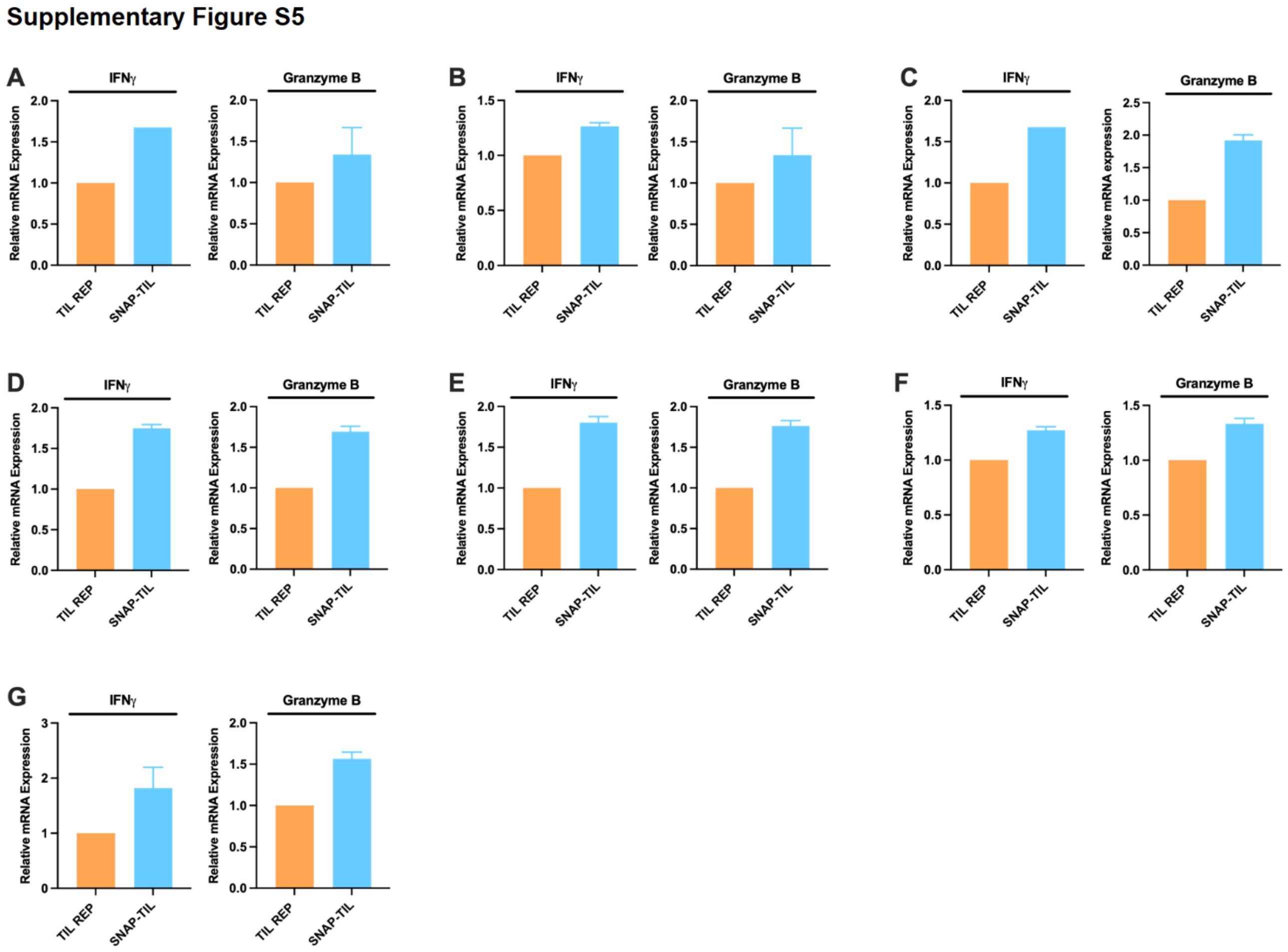

**Supplementary Figure S6**

IFNγ and granzyme B levels in TILs co-cultured with autologous *ex-vivo* tumoroids measured by MSD assay. (A) CRC, (B) melanoma, (C) uveal melanoma, (D) NSCLC, (E-I) PDAC. Each bar represents the mean ± SD, n = 2 -3, $^{**}p < 0.01$, $^{***}p < 0.001$, ordinary one-way ANOVA with Dunnett's post-hoc test.

**Supplementary Figure S6**

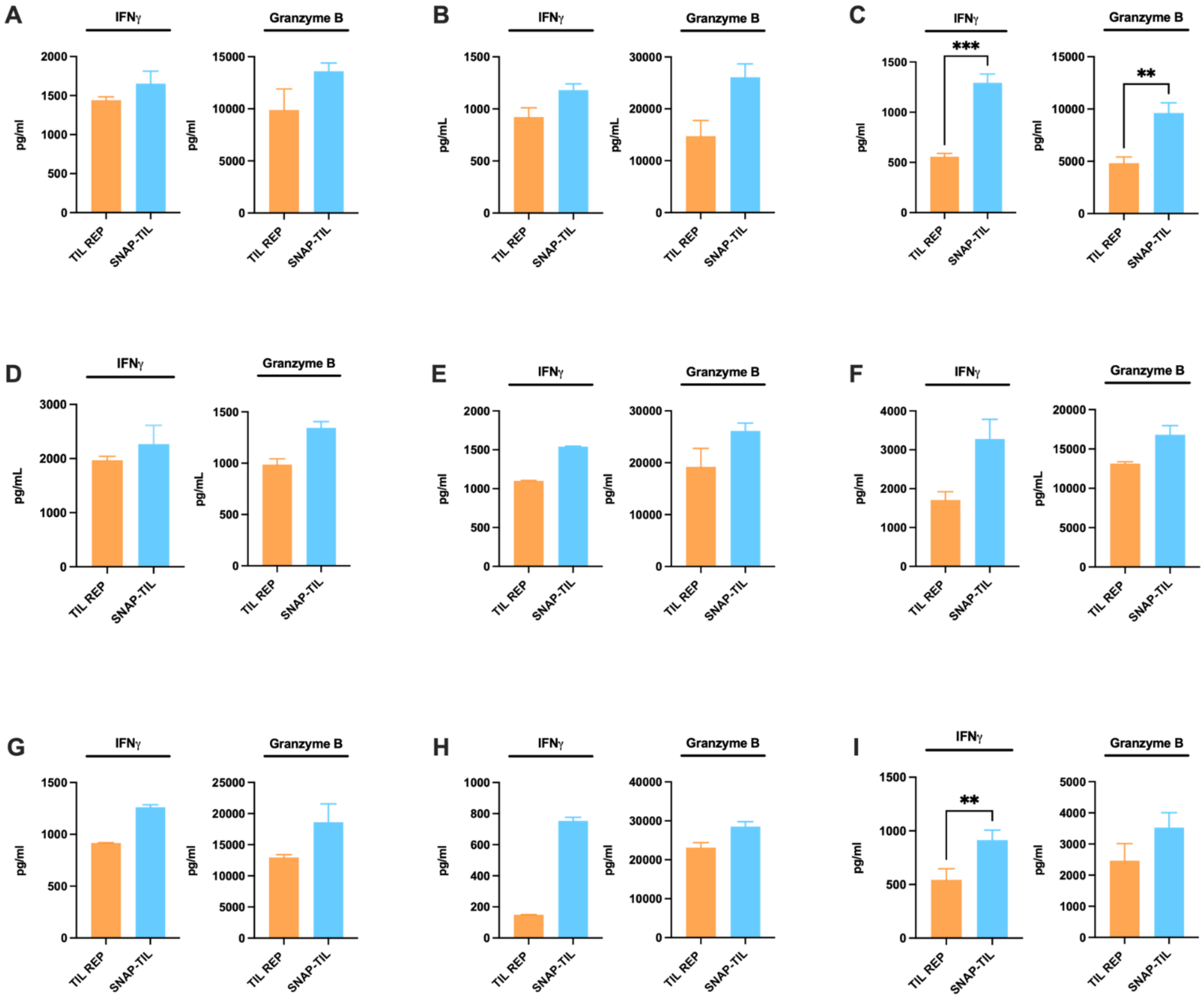

**Supplementary Figure S7**

Female *hIL2*-NOG mice were subcutaneously implanted with propagated melanoma or PDAC cells. Once tumors reached around 200 mm$^3$, indicated by a dotted line, autologous SNAP-TIL, TIL REP, or vehicle was administered by retroorbital injection. Body weight was measured weekly during the study and plotted as the mean ± SEM, n = 10 – 14 mice per group. (A) melanoma or (B) PDAC PDX models.

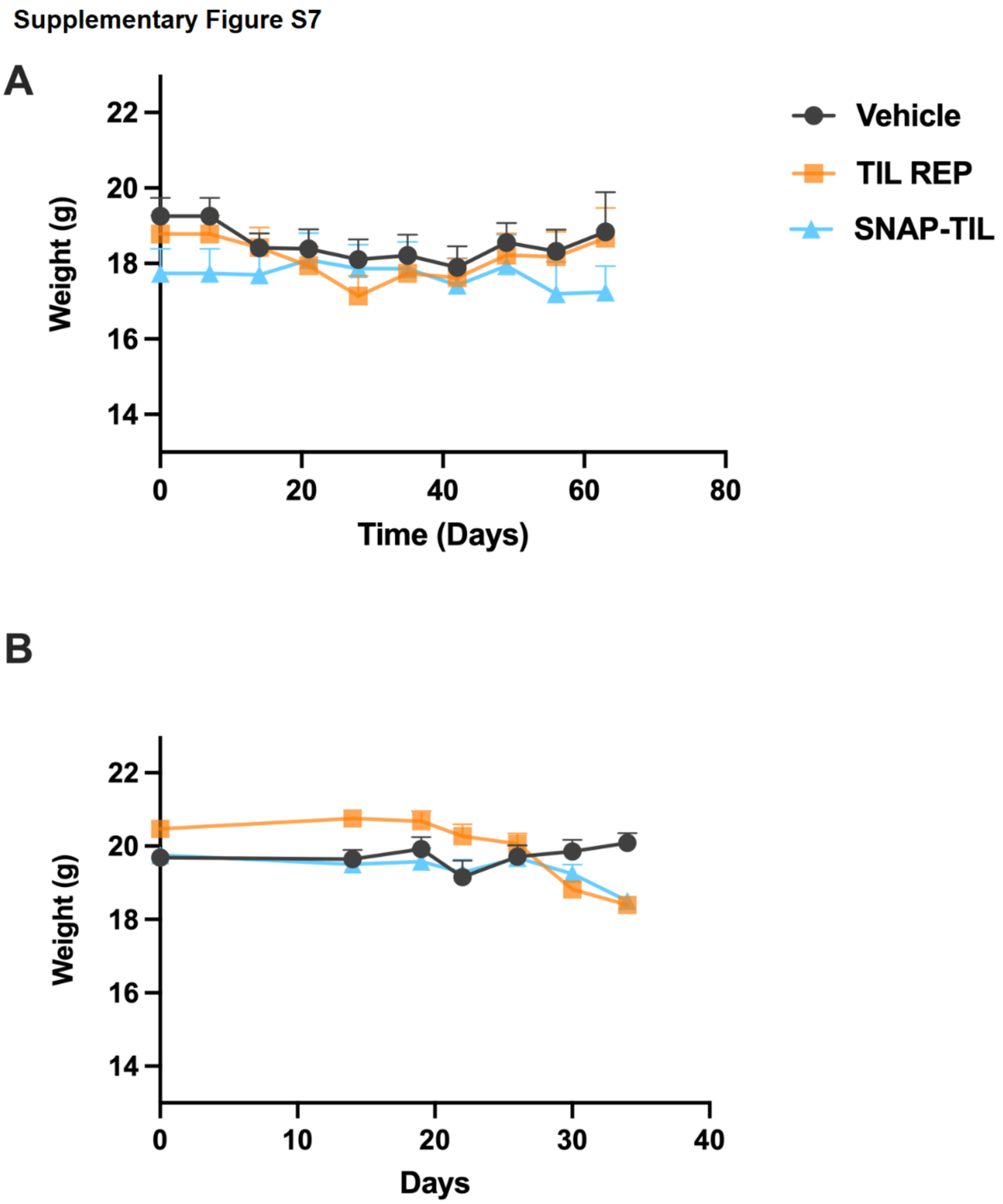
