## Supplementary table for "Preclinical Proof of Concept for a personalized SNAP^TM^-TIL (Specific Neo-Antigen Peptides -TIL) therapy platform"

**Supplementary Table S1. Antibodies used for flow cytometry analysis.**

| Antibody | Clone | Company | Catalog Number |
|---|---|---|---|
| FITC anti-human CD326 (EpCAM) | 9C4 | Biolegend | 324203 |
| FITC anti-human CD45 | 2D1 | Biolegend | 368507 |
| PE anti-human CD45 | 2D1 | Biolegend | 368509 |
| APC anti-human CD45RO | UCHL1 | Biolegend | 983102 |
| PE/Cyanine7 anti-human CD45 | 2D1 | Biolegend | 368531 |
| PerCP/Cyanine5.5 anti-human CD45 | 2D1 | Biolegend | 368503 |
| FITC anti-human CD4 | A161A1 | Biolegend | 357406 |
| PE anti-human CD8 | SK1 | Biolegend | 344706 |
| FITC anti-human CD19 | 4G7 | Biolegend | 392508 |
| PE/Cyanine7 anti-human CD3 | HIT3a | Biolegend | 300316 |
| PE anti-human CD56 (NCAM) | 5 1H11 | Biolegend | 362508 |
| PerCP/Cyanine5.5 anti-human CD28 | CD28.2 | Biolegend | 302921 |
| APC anti-human CD27 | M-T271 | Biolegend | 356409 |
| PE/Cyanine7 anti-human CD45RA | HI100 | Biolegend | 304125 |
| APC anti-human CD45RO | UCHL1 | Biolegend | 983102 |
| FITC anti-human CD62L | DREG-56 | Biolegend | 304804 |
| APC anti-human CD197 (CCR7) | G043H7 | Biolegend | 353213 |
| PE anti-human FOXP3 | 206D | Biolegend | 320107 |
| Annexin v/7-AAD | NA | Biolegend | 640922 |
| PE anti-human CD95 | DX2 | Biolegend | 305607 |
| FITC anti-human IL-7Ra | A019D5 | Biolegend | 351311 |

**Supplementary Table S2. Primer list for qPCR.**

| Gene Symbol | Forward Primer (5'-3') | Reverse Primer (3'-5') |
|---|---|---|
| *TIGIT* | CGTGAACGATACAGGGGAGT | GCAATGGAATCTGGAACCTG |
| *PD1* | ACCTGGGTGTTGGGAGGGCA | GGAGTGGATAGGCCACGGCG |
| *TIM3 (HAVCR2)* | AGGGGACATGGCCCAGCAGA | GCCAGCCCAGCACAGATCCC |
| *Hu Lag-3* | CTAGCCCAGGTGCCCAACGC | GCCTGCGGAGGGTGAATCCC |
| *IFNG* | TGACCAGAGCATCCAAAAGA | CTCTTCGACCTCGAAACAGC |
| *GZMB* | TGCAGGAAGATCGAAAGTGCG | GAGGCATGCCATTGTTTCGTC |
| *PRF1* | ACCAGCAATGTGCATGTGTCTGTG | GAAGGAGGCCGTCATCTTGTGCTT |

*TIGIT*, T cell immunoreceptor with Ig and ITM domains; *PD1*, programmed cell death; *TIM3*, T-cell immunoglobin and mucin domain 3, IFNG, interferon-gamma; *GZMB*, granzyme B; *PRF1,* perforin 1

**Supplementary Table S3.** Percent of CD4, CD8, or IFNγ-positive cells per PDAC PDX tumor volume between REP-TIL and snapTIL treated mice. Cell counts measured by IHC (cell/$mm^2$).

| Sample ID | TIL received | Initial tumor volume ($mm^3$) | End tumor volume ($mm^3$) | % CD4+ | % CD8+ | % IFNγ + |
|---|---|---|---|---|---|---|
| I2-03 | REP TIL | 192.196 | 552.202 | 1.89 | 3.47 | 1.06 |
| I2-13 | REP TIL | 192.196 | 567.176 | 1.60 | 2.03 | 1.29 |
| I3-43 | REP TIL | 192.196 | 788.170 | 0.60 | 2.84 | 0.86 |
| S3-13 | snapTIL | 192.196 | 447.306 | 2.51 | 3.81 | 2.98 |
| S2-11 | snapTIL | 192.196 | 298.223 | 4.86 | 5.35 | 4.67 |
| S1-30 | snapTIL | 192.196 | 322.464 | 3.78 | 5.82 | 3.82 |
